# Evolutionary adaptation under climate change: *Aedes* sp. demonstrates potential to adapt to warming

**DOI:** 10.1101/2024.08.23.609454

**Authors:** Lisa I. Couper, Tristram O. Dodge, James A. Hemker, Bernard Y. Kim, Moi Exposito-Alonso, Rachel B. Brem, Erin A. Mordecai, Mark C. Bitter

## Abstract

Climate warming is expected to shift the distributions of mosquitoes and mosquito-borne diseases, facilitating expansions at cool range edges and contractions at warm range edges. However, whether mosquito populations could maintain their warm edges through evolutionary adaptation remains unknown. Here, we investigate the potential for thermal adaptation in *Aedes sierrensis*, a congener of the major disease vector species that experiences large thermal gradients in its native range, by assaying tolerance to prolonged and acute heat exposure, and its genetic basis in a diverse, field-derived population. We found pervasive evidence of heritable genetic variation in acute heat tolerance, which phenotypically trades off with tolerance to prolonged heat exposure. A simple evolutionary model based on our data shows that the estimated maximum rate of evolutionary adaptation in mosquito heat tolerance typically exceeds that of projected climate warming under idealized conditions. Our findings indicate that natural mosquito populations may have the potential to track projected warming via genetic adaptation. Prior climate-based projections may thus underestimate the range of mosquito and mosquito-borne disease distributions under future climate conditions.

**Significance Statement:** Global change may have profound impacts on the distribution of mosquito-borne diseases, which collectively cause nearly one million deaths each year. Accurately predicting these impacts is critical for disease control preparedness, and will depend, in part, on whether mosquitoes can adapt to warming—a key open question. Using experimental and genomic data from a relative of major vector species that already experiences a wide thermal gradient, we find that natural mosquito populations have high levels of genetically-based variation in heat tolerance that could enable adaptation on pace with warming. Incorporating the potential for adaptive responses may therefore be necessary for accurate predictions of mosquito-borne disease distributions under warming, which is critical for preparing mosquito control interventions.

## Introduction

Climate warming is expected to alter the global distributions of mosquitoes that transmit pathogens, disrupting existing vector control measures and changing the landscape of disease risk (1, 2). Mosquito species ranges are constrained by their thermal limits, which dictate the suitable temperatures over which they can survive, develop, and reproduce. Consequently, mosquito ranges are predicted to shift with warming, expanding polewards and towards higher altitudes as temperatures become newly suitable at current cool range edges, and contracting at current warm edges as temperatures become newly prohibitive (3–5). For mosquitoes that transmit diseases, including dengue, malaria, and West Nile virus, which collectively cause nearly one million deaths annually (6), this process is already underway, as warming-related range expansions have been observed for several species of *Anopheles* (7–9), *Aedes* (10, 11), and *Culex* (12, 13) mosquitoes. However, most mosquito and vector-borne disease models project that mosquito distributions will also contract at warm edges as temperatures begin to exceed their upper physiological limits. Whether such warming-driven contractions will actually occur, or whether evolutionary adaptation may enable populations to maintain their warm range edges as temperatures increase, is unknown (14–16). Determining thermal adaptive potential for mosquito species will both augment our understanding of species responses to climate change and inform management strategies for controlling disease spread as climate change progresses.

To persist near current warm edges, mosquitoes may need to rapidly adapt to temperatures beyond their current upper thermal limits. Several common properties of mosquitoes indicate that rapid adaptation is feasible, including short generation times, large population sizes, and steep declines in fitness above their thermal optima (reviewed in (15)). However, the extent of variation and heritability in heat tolerance—fundamental components of thermal adaptive potential—remain poorly understood for most mosquito species. Several prior studies have found phenotypic variation in heat tolerance for populations of *Aedes* (17), *Anopheles* (18), *Culex* (19–22), and *Wyeomyia* (23) species when assessed under constant temperature exposures in the lab. While this phenotypic variation putatively reflects heritable variation for thermal tolerance as the studies typically controlled for direct environmental effects (*i.e.,* by using common garden experimental designs), the genomic basis and extent of genetic variation in heat tolerance were not directly investigated. Further, while prior studies have investigated mosquito thermal performance for several life history traits (*e.g.*, larval development rates, pupal survival, adult lifespan)(17–19, 21, 24), the extent to which thermal tolerance at juvenile stages is predictive of tolerance at later developmental stages, and whether such cross-stage tolerances are genetically correlated, remains unknown. Finally, several additional studies have found strong direct evidence for heritable variation in response to acute heat shock in adult *Ae. aegypti* (24, 25), but the underlying mechanisms and genetic basis of this variation were not identified.

The pace at which mosquitoes adaptively track warming temperatures may in large part hinge upon the underlying genetic architecture of thermal tolerance, including the number of independent loci underpinning phenotypic variation in tolerance and the distribution of these loci throughout the genome (26, 27). Across a diverse range of taxa, traits involved in climate adaptation typically exhibit a polygenic basis, whereby hundreds to thousands of genes underpin adaptive phenotypes (28–33). These adaptive loci have often been shown to cluster within chromosomal inversions, a form of structural mutation in which segments of DNA are broken off and become reattached in the reverse orientation. Suppressed recombination within inversion breakpoints can then facilitate the co-segregation of adaptive alleles and augment their spread within populations (34, 35). In *Anopheles* spp., inversions have been found to underscore adaptive traits including desiccation resistance, larval thermal tolerance, insecticide resistance, host preference, and ecotype formation (36–42). Inversions have also been found to be abundant in *Aedes* spp., however their role in climate adaptation remains largely unknown (43–45). Overall, the underlying genetic architecture of heat tolerance, including the role of chromosomal inversions, remains poorly understood for most mosquito species, hindering efforts to predict the capacity for adaptive evolution on pace with climate warming.

Here, we investigated the potential for evolutionary adaptation in heat tolerance using the western tree hole mosquito, *Aedes sierrensis*, as a model system. In addition to being a major pest species and vector of canine heartworm in western North America, as well as a congener of the major human disease vector species (*i.e., Ae. aegypti* and *Ae. albopictus*), *Ae. sierrensis* is abundant, distributed across a wide climate gradient (ranging from southern California to British Columbia)(46), and easy to identify, sample, and manipulate in the lab. Leveraging these properties, we sought to answer the following specific research questions: (i) How much standing variation in heat tolerance exists in natural mosquito populations? (ii) How does prolonged heat exposure at the larval stage impact acute heat tolerance at the adult stage? (iii) What is the genetic architecture of these short- and long-term heat tolerance traits? (iv) Could standing variation in thermal tolerance enable natural populations to adapt on pace with climate warming, altering projections of future range shifts?

To answer these questions, we conducted a thermal selection experiment in which we reared a genetically diverse starting population, derived from the center of the *Ae. sierrensis* range, at high temperatures (30°C) that approximately capture the upper thermal limits for prolonged larval survival, or control (22°C) temperatures that approximately capture the maximum temperature experienced by this population during the larval activity season. Using surviving individuals from both temperature conditions, we assayed acute heat tolerance using a thermal knockdown assay, and conducted a genome-wide association analysis of acute and prolonged heat tolerance. We found large phenotypic variation in acute heat tolerance within the study population, and a putative trade-off in heat tolerance to prolonged versus acute exposure, whereby individuals reared under high temperatures during larval development had significantly lower acute heat tolerance as adults. Our genomic analysis revealed a polygenic architecture of both heat tolerance traits, and a putative role of chromosomal inversions underpinning thermal adaptation within the species. Lastly, using parameter estimates derived from our experimental and genomic data, we estimated that the maximum rate of evolutionary adaptation in larval heat tolerance typically exceeds that of projected climate warming under simplified conditions. This finding suggests that natural populations may harbor the potential to adapt on pace with warming, and that prior climate-based projections that do not incorporate adaptation may underestimate the ranges of mosquitoes and mosquito-borne disease transmission under future climate conditions.

## Results

### Extent of variation in acute heat tolerance

Our thermal selection experiment was conducted on a large, diverse starting population of *Ae. sierrensis* collected from tree hole habitats across Solano County, CA (mean **π** = 0.0015; see Supplemental Figures S1-S2 and Supplemental Table S4 for additional population diversity metrics). From this starting population, we found large individual-level variation in acute heat tolerance. Specifically, we reared field-collected individuals at common temperatures (22°C) for two generations, then implemented a selection experiment design in which F_3_ larvae were either maintained at control temperatures of 22°C—approximately the maximum temperature this population currently experiences during the spring when larvae are developing—or high temperatures of 30°C—approximately the upper thermal limits for prolonged larval survival (17)(n = 790 total control larvae, 1,943 heat-selected larvae; see Supplemental Table S1 for sample numbers per experimental round). Surviving individuals from both treatments were returned to control conditions (22°C) at pupation and reared to adulthood. These temperature conditions generated substantial differences in larval survival between treatments: survival dropped from 57.8% in the control group to 24.2% in the heat-selected group (across all experimental rounds, see Supplemental Table S1 for rates per round). Although individuals from both groups experienced the same temperature as pupae (22°C), individuals in the heat-selected group retained lower survival rates at this life stage compared to the control group (74.3% and 93.2% pupal to adult survival, respectively). This resulted in overall survival rates from larvae to adulthood of 18.0% in the heat-selected group and 53.5% in the control group.

Using all individuals that survived to adulthood from either treatment (n = 122 control, 105 heat-selected individuals), we conducted a thermal knockdown assay—the time to loss of motor function in a warm water bath—a frequently used proxy for acute heat tolerance whereby longer knockdown times indicate greater heat tolerance (24, 25, 47, 48). Our results indicate large individual-level variation in acute heat tolerance (Figure 2), with adults from the control group ranging in knockdown times from 32.7 to 67.6 minutes (median: 48.8 minutes) and those from the heat-selected group ranging from 19.8 to 64.8 minutes (median: 46.2 minutes)(t = 2.65, p < 0.01; differences between groups discussed further below). The variance in knockdown times was marginally larger for the heat-selected group (83.8 and 76.9 for males and females, respectively) than for the control group (56.9, 55.7)(F = 0.071, p = 0.06 for both sexes combined; F = 0.68, p = 0.15 for males only; F = 0.72, p = 0.23 for females only).

**Figure 1.**
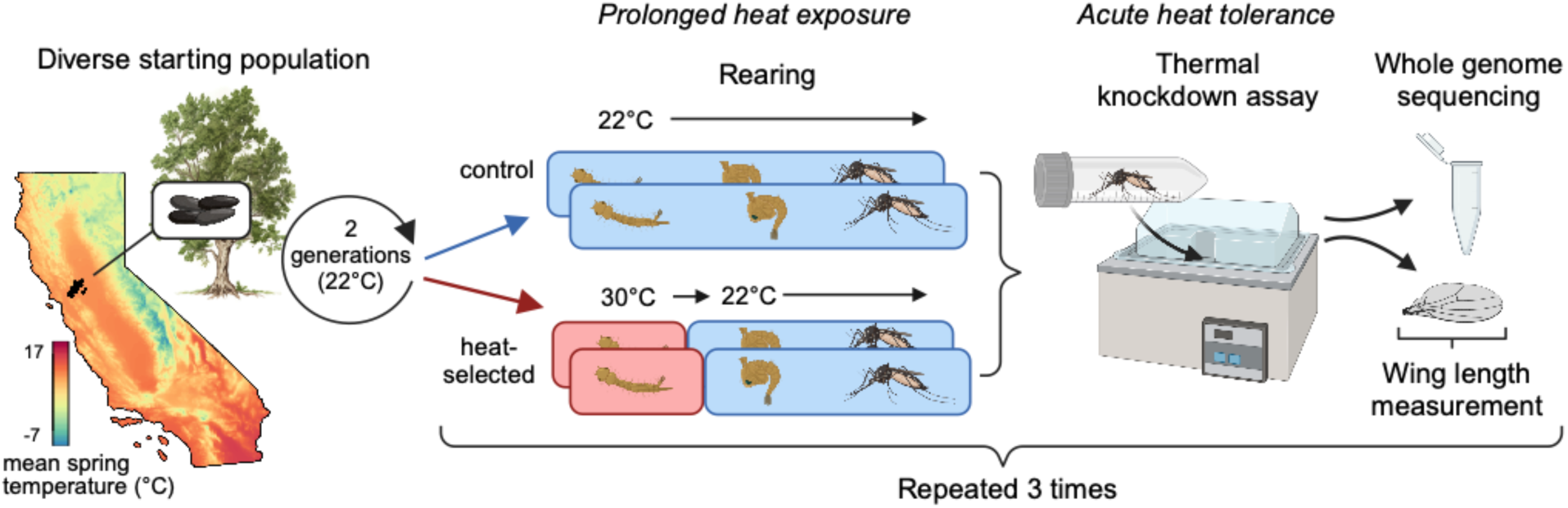
Assessing chronic and acute heat tolerance in a genetically diverse field-derived population of mosquitoes. A diverse starting population was obtained from tree hole habitats across Solano County, CA., with the original sampling locations denoted on the map. Map colors denote variation in average daily temperatures in the spring—the larval activity period. All individuals were reared under lab conditions for two generations, and the resulting F_3_ eggs were used in the experiment. Eggs were hatched at 22°C, and 24-h larvae were randomly designated into replicated control or heat-selected groups. Individuals were reared at 22°C (control) or 30°C (heat-selected) as larvae. All individuals were maintained at 22°C at the pupal and adult life stages. Acute heat tolerance was assayed via thermal knockdown on individuals 48-72h after eclosion. All individuals were then preserved for DNA sequencing and body size approximation. The full experiment was conducted 3 times for biological replication.

**Figure 2.**
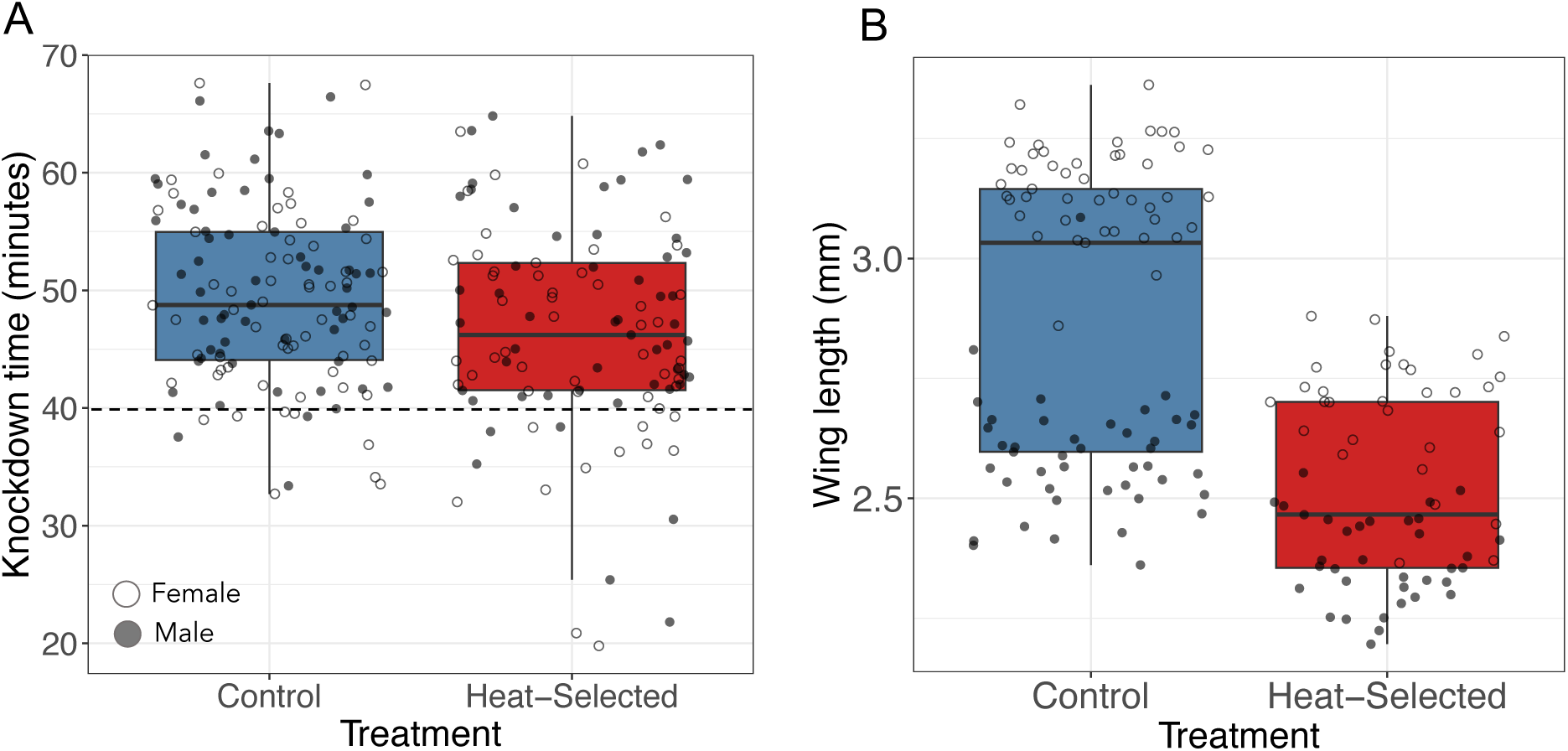
Acute adult thermal tolerance and wing length are reduced in individuals that experienced larval heat selection. Variation in (A) thermal knockdown time (a metric of upper heat tolerance) and (B) wing length (a metric of body size) measured in control (blue) and heat-selected (red) *Ae. sierrensis* adults. Each point denotes the knockdown time or wing length of a single assayed individual. Heat-selected individuals knocked down significantly quicker and had smaller body sizes than control individuals (p = 0.03, < 0.001). Open and filled circles denote females and males, respectively. Points are jittered to aid in visualization. The dashed horizontal line at 40 minutes on the left plot denotes the time the water bath reached the final set temperature of 38°C. Points below this line thus denote individuals that knocked down at a lower temperature.

### Impact of prolonged larval heat exposure on acute adult heat tolerance

We found that mosquitoes that underwent heat-selection as larvae had significantly shorter knockdown times as adults than the control group (LMM, t = −2.15, p = 0.03; Figure 2A). Specifically, heat-selected larvae knocked down, on average, 3.6 minutes earlier than control larvae (46.1 ± 8.9, 49.7 ± 7.4 minutes for heat-selected and control larvae, respectively). To identify potential mechanisms underlying variation in acute heat tolerance, we measured the wing length—a validated proxy for overall mosquito body size—of each individual used in the thermal knockdown assay (49). We found that adults that underwent heat-selection as larvae had significantly smaller wing lengths than those from the control group (LMM, t = −16.21, p < 0.001, Figure 2B). Specifically, the average wing length for the heat-selected group was 0.31 mm smaller than the control group (2.51 ± 0.19 and 2.86 ± 0.30mm, respectively; Supplemental Table S2). However, while wing lengths differed between treatment groups, wing length itself was not a significant predictor of knockdown time (LMM, t = −1.17 p = 0.24; Supplemental Table S3). Wing lengths also varied significantly by sex (LMM, t = −22.26, p <0.001, Figure 2), with the average female wing length being 0.47 mm larger than that of males (2.96 ± 0.25 and 2.49 ± 0.15 mm, respectively; Supplemental Table S2).

### Genomic architecture of prolonged and acute heat tolerance

We conducted whole genome sequencing on all 227 adult *Ae. sierrensis* adults used in the experiment, obtaining an average of 95 million reads and a sequencing depth of ∼10X per individual (see Supplemental Tables S4-S5 for per-sample summary statistics). We aligned these reads to our de novo *Ae. sierrensis* reference genome assembly (1.183 Gb; available at NCBI BioProject ID: PRJNA1119052 upon manuscript acceptance), and, after quality control and variant filtering, identified 583,889 single nucleotide polymorphisms (SNPs) segregating in our study population. We leveraged these polymorphic sites to identify genomic regions associated with prolonged and acute heat tolerance. For prolonged heat tolerance, we used an F_ST_ outlier approach to identify SNPs with elevated differentiation between control and heat-selected groups and, in addition, a case-control genome-wide association (GWA) between control and heat-selected individuals (see Methods*: Identifying genetic variants associated with heat tolerance*). We leveraged the overlap in these approaches to more robustly identify genes underpinning differences in survival between treatments. For acute heat tolerance, we implemented a standard genome-wide association (GWA) analysis using adult knockdown time as the dependent variable.

We identified hundreds of candidate SNPs, distributed across all three chromosomes, associated with thermal tolerance. Specifically, we identified 351 and 113 outlier SNPs associated with prolonged larval heat tolerance via the F_st_ outlier (q < 0.05 and F_st_ ≥ 0.05) and case-control GWA (FDR-corrected p < 0.01), respectively (Figure 3A, Supplemental Figure S3). Both approaches produced clusters of significant SNPs in distinct regions of chromosome one, two, and three (Figure 3A, Supplemental Figure S3), putatively indicating a set of tightly linked genes, or larger structural variants, driving the signal in these regions. The GWA of knockdown time yielded 120 candidate SNPs, but with a much more diffuse distribution across the genome (Figure 3B). As expected, across all candidate SNPs, we quantified systematic allele frequency differences between treatment groups (Figure 3D-E). That is, we found significantly larger differences in candidate SNP frequency between control and heat-selected individuals, or between individuals with high (upper 50% of phenotypic distribution) and low (bottom 50%) knockdown times, relative to a set of matched controls (see Methods: *Estimating allele frequency shifts*)(Figure 3D-E, Supplemental Figure S4). Together, these findings indicate that heat tolerance—to both prolonged and acute heat stress—results from allele frequency shifts at SNPs located throughout the genome, ultimately suggesting that polygenic adaptation may underpin population persistence under warming.

**Figure 3:**
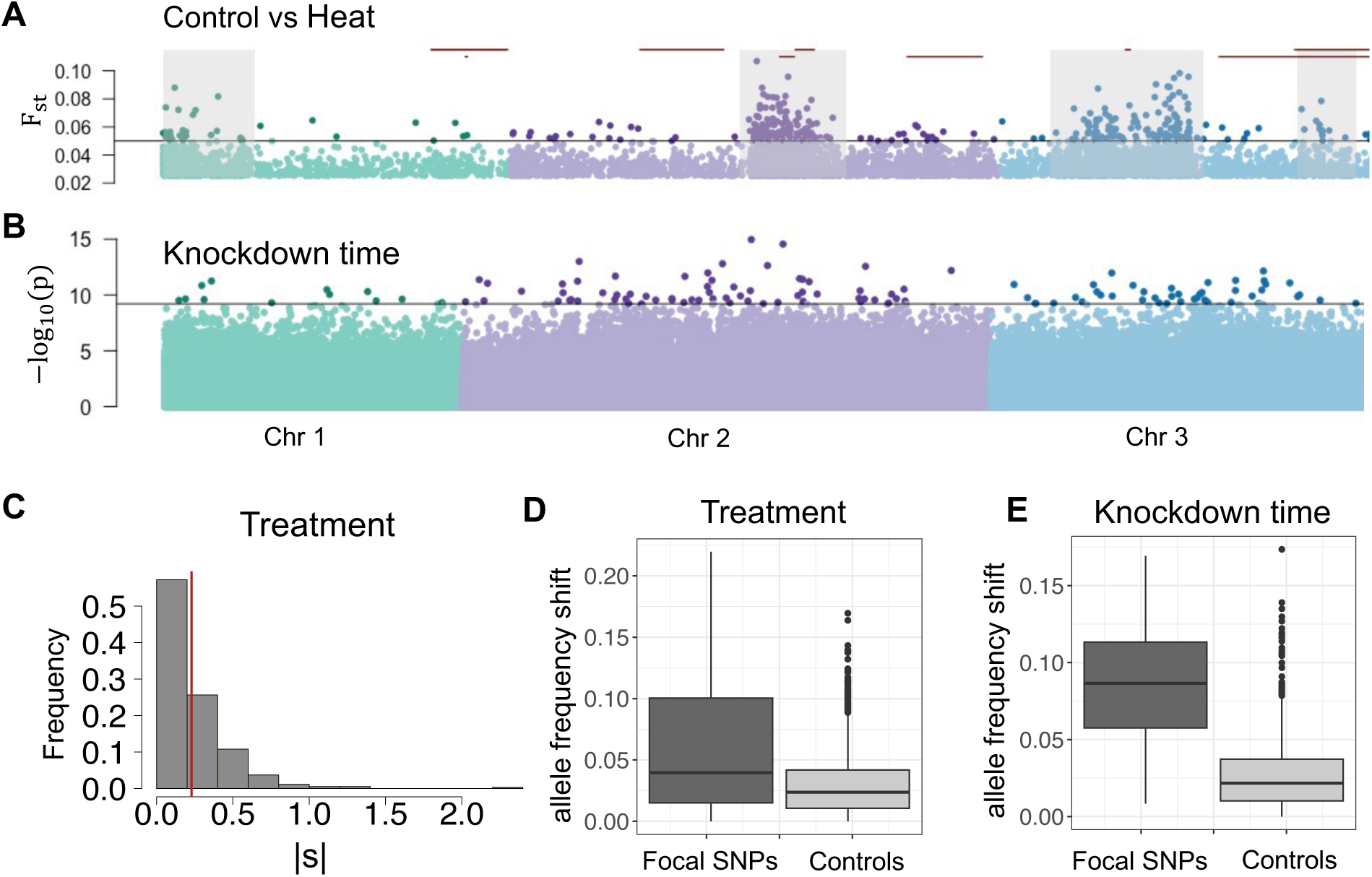
The genomic architecture of thermal tolerance. (A) Genomic position of candidate SNPs significantly associated with larval heat treatment (a measure of prolonged heat tolerance). The black horizontal line indicates the threshold for significance as candidate SNPs (*i.e.*, q < 0.05 and F_ST_ > 0.05). Gray shaded rectangles denote regions with an enrichment of SNPs differentiating treatment groups, relative to the genome-wide average. Red horizontal lines denote the location of chromosomal inversions significantly differentiated in frequency between control and heat-selected larvae. (B) Genomic position of candidate SNPs significantly associated with knockdown time (a measured of acute heat tolerance). Here, the black horizontal line indicates the threshold for significance as candidate SNPs based on FDR-corrected p < 0.001. (C) Distribution of selection coefficients, |s|, for candidate SNPs associated with prolonged heat tolerance at the larval stage. The red vertical line denotes the mean. (D) Difference in allele frequency distributions for focal SNPs (dark gray) versus their matched controls (light gray) for the larval heat treatment. Here, the focal SNPs from the F_ST_ and case-control GWA approaches are shown together. (E) Difference in allele frequency distribution for focal SNPs (dark gray) versus their matched control (light gray) for adult thermal knockdown. The black line in each boxplot denotes the median allele frequency difference. See Supplemental Figure S4 for allele frequency shifts based on starting minor allele frequency.

The large chromosomal regions enriched in SNPs associated with prolonged thermal tolerance (Figure 3A) indicated that larger structural variants, or regions of co-adapted gene complexes, may underpin phenotypic variation in this trait. To investigate this, we used genomic data to assess whether chromosomal inversions, which are known to play a pronounced role in climate adaptation in ectotherms including mosquito species (37, 50), segregate in our study population and potentially differentiate individuals based on their thermal tolerance. Using consensus results from four short-read structural variant callers, we identified 444 inversions segregating in our focal populations (after filtering based on size between 1-200 Mb and frequency >5%). These were indeed enriched in the specific chromosomal regions where we observed a dramatic elevation in the number of SNPs associated with prolonged, larval heat tolerance (Figure 3A, Supplemental Methods). That is, 59% (n = 261) of the inversions occurred within these regions of interest, which span approximately 31% of the genome. We identified a total of nine inversions that were significantly differentiated in frequency between control and heat-selected individuals, five of which fell within these regions of interest (Figure 3A, Supplemental Figure S5, Supplemental Table S6). We further identified five inversions that were significantly differentiated between individuals from the top 25% and bottom 25% of knockdown times, after controlling for treatment and sex (Supplemental Table S6). While the patterns observed here, in combination with mounting research in congeners (37, 51, 52), suggests inversions may underpin climate adaptation in a range of mosquito species, inferences from short-read sequencing data are limited and future research leveraging long-read sequencing data to validate and resolve the particular inversions segregating and driving phenotypic differentiation is warranted.

We next generated a list of genes putatively underlying outlier SNP differentiation. Specifically, by identifying all genes within 50 kb of each focal SNP, we identified 293, 105, and 114 annotated genes for the aforementioned three approaches, respectively (*i.e.*, F_st_ and GWA between control and heat-selected individuals, and GWA on knockdown time) out of a total of 30,554 predicted genes across the genome (Supplemental Figure S6). We found that the number of overlapping candidate genes identified by the two approaches capturing prolonged heat tolerance (n = 13) was significantly greater than expected by chance (95% confidence interval from expectation: 1.30 - 1.51, p < 0.01)(Supplemental Figure S6), supporting the robustness of these two methods at capturing similar candidate genes from the same phenotypic data. These overlapping genes mapped to genes previously associated with heat or environmental stress responses in other ectotherm species including histone H3 (involved in heat shock memory (53, 54)), profilin (an actin-binding protein that may function as a molecular chaperone (55, 56)), and cytochrome P450 (involved in thermal stress responses in a wide range of taxa (57–59))(Supplemental Table S7). The number of overlapping genes between the prolonged and acute heat tolerance approaches (n = 3, 3) was also significantly greater than expected by chance (95% CI: 1.30 - 1.50, 0.48 - 0.62, p < 0.01 for both)(Supplemental Figure S6), potentially suggesting shared genetic pathways between prolonged and acute heat tolerance phenotypes.

### Investigating potential to adapt on pace with warming

We investigated whether the level of standing variation in heat tolerance observed here could fuel adaptation on pace with climate warming using a simple evolutionary rescue model parameterized by our experimental data. Specifically, we estimated the maximum rate of evolutionary adaptation in prolonged larval heat tolerance, and compared this to rates of change in mean daily temperature during the larval activity period (typically January - April), projected under a moderate warming scenario (RCP 4.5). We focused on larval heat tolerance as our recent investigation of thermal tolerance across life stages in this species indicated larval survival may be the bottleneck to thermal adaptation (17). We note several simplifying assumptions that may limit the accuracy of our predictions including the absence of phenotypic plasticity and/or impacts of concurrent abiotic and biotic stressors in the model; the focus on a single trait underlying adaptation (*i.e.*, no constraints or trade-offs); the assumption of constant heritability and phenotypic variance over time and uniform genomic variance across space (*i.e.,* range edge populations are assumed to have similar genetic variation as central populations); and the use of selection strength estimated as the difference in survival between our experimentally-imposed temperature treatments (*i.e*., 30 vs 22°C), which was guided by, but not specific to, the shift in temperatures that natural populations may experience (see Methods: *Estimating adaptive potential*). For these reasons, we use this modeling approach to estimate the potential parameter space of evolutionary adaptation under idealized conditions, rather than to obtain a precise estimate of adaptation expected under natural settings. Using this approach, we found that estimated rates of adaptation exceed rates of warming under most of our experimentally-estimated or literature-derived parameter values (Figure 4, Supplemental Figure S7). Specifically, the maximum estimated rate of evolutionary change derived from our experimental and genomic data (*i.e.*, point estimates for selection strength, heritability, and phenotypic variation) and previously estimated rates of maximum mosquito population growth (r_max_) ranged from 0.033 – 0.051 °C/year (for r_max_= 0.15 and 0.35, respectively), exceeding projected rates of warming in mean spring temperatures across the southern portion of the *Ae. sierrensis* distribution (0.026 °C/year). We consider rates of warming in mean spring temperatures across the southern *Ae. sierrensis* distribution projected under RCP 4.5 to be the most ecologically relevant metric of warming, but we also consider alternative metrics including the projected rate of change in maximum daily spring temperatures and mean annual temperatures across this same extent under RCP 4.5 (0.040, 0.015 °C/year, respectively), the projected rate of change in annual mean temperatures across the southeastern U.S. under RCP 4.5 and RCP 8.5 (0.031, 0.035 °C/year, respectively), and recently observed rates of warming in annual mean temperature across North America (0.027 °C/year) (Supplemental Table S9, Supplemental Figure S7). Across these metrics of warming, our results provide empirical support for the potential for evolutionary adaptation in this population in response to climate warming

**Figure 4.**
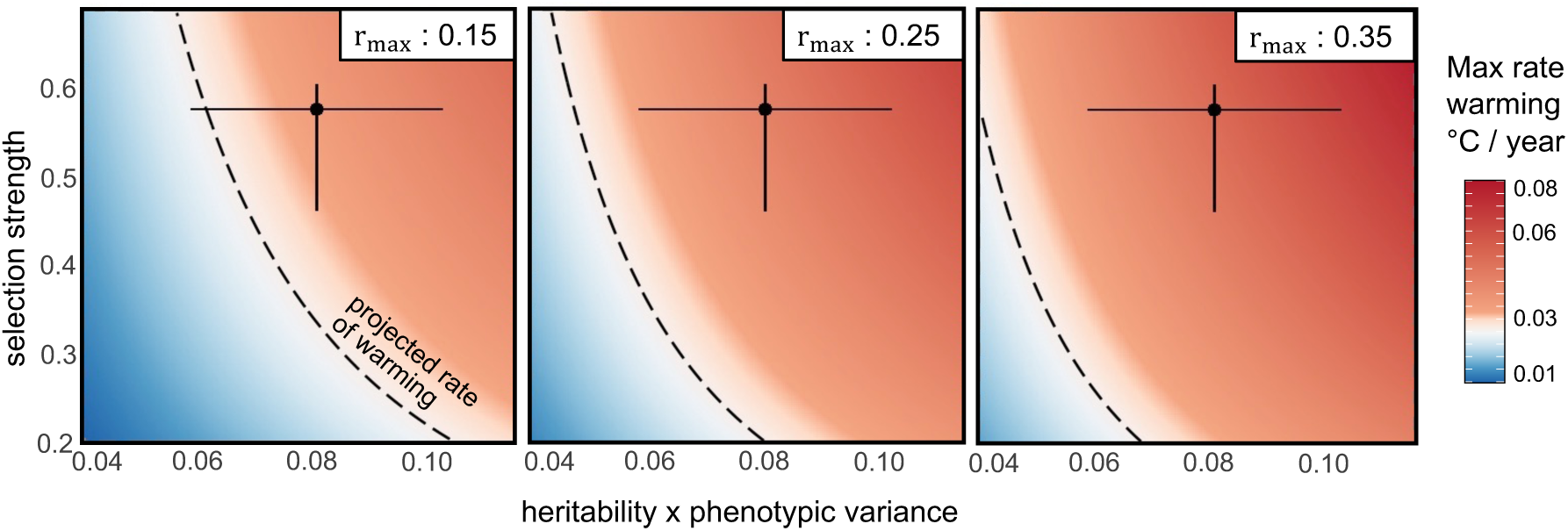
Rates of evolutionary adaptation typically exceed projected rates of climate warming. Colors denote the maximum rate of warming (°C/year) to which populations could adapt (equivalent to the model-estimated maximum rate of evolutionary change). The x-axis denotes potential values for the product of heritability (h^2^) and phenotypic variance (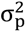), and the y-axis denotes potential values for selection strength (γ). The black circle on each plot denotes the point estimate for these three parameter values from our experimental and genomic data and the error bars capture the range of these parameters made under varying model assumptions (see Methods: *Estimating Adaptive Potential*). The isoline denote the rates of warming in daily mean temperature during the spring across the southern portion of *Ae. sierrensis* distribution (0.026°C/year). Error bars and point estimates to the right of the isoline reflect scenarios under which the population’s estimated maximum rate of evolutionary adaptation exceeds the projected rate of warming. The three panels span previously estimated rates of maximum mosquito population growth rates (r_max_ = 0.15, 0.25, and 0.35 for the left, center, and right panels, respectively).

## Discussion

Nearly all climate-based projections of future mosquito and mosquito-borne disease distributions assume that mosquitoes will migrate to track their current niches rather than evolve in response to temperature change (15, 24, 60). We examined the potential for thermal adaptation using a selection experiment conducted on the western tree hole mosquito, *Aedes sierrensis*, a congener of major disease vector species, *Ae. aegypti* and *Ae. albopictus*. Our results suggest that evolutionary adaptation is likely a viable and important component of mosquito responses to climate warming.

### Large within-population variation in acute heat tolerance

We found a high level of standing phenotypic variation in heat tolerance within a single, field-derived mosquito population, underpinned by several hundred genes and an abundance of chromosomal inversions. In particular, when exposed to acute high temperature stress, we found that the time to loss of motor function (*i.e.,* ‘knockdown time’) varied from 32.7 to 67.6 minutes between individuals from a single starting population (here, the control group), despite having experienced the same thermal environment for the prior two generations (*i.e.,* 22°C). This variation in acute heat tolerance was even larger—ranging from 19.8 to 64.8 minutes—in individuals that were exposed to prolonged heat stress as larvae (*i.e.,* treatment group, 30°C). These findings match recent observations of large and heritable variation in acute heat tolerance in related *Aedes* species (24, 25). As genetically-based trait variation is critical for adapting to changing conditions (61, 62), our results suggest that *Ae. sierrensis* may harbor standing variation in heat tolerance that could enable adaptation to aspects of climate warming.

### Prolonged heat exposure as larvae led to lower acute heat tolerance at the adult stage

Our selection experiment also revealed potential costs or trade-offs between prolonged heat exposure during rearing and acute heat tolerance in subsequent life stages. In particular, we found that individuals reared at 30°C (‘heat-selected’) as larvae had significantly lower acute heat tolerance as adults (as evidenced by quicker knockdown times) than those reared at 22°C (‘control’). This finding appears to contrast with prior empirical and theoretical work in thermal biology finding that exposure to high temperatures at early life-stages leads to acclimation and higher heat tolerance in adulthood for ectotherms (63–70), including in related *Aedes* species (71). However, our result may be explained by well-supported mechanisms including variation in adult body size resulting from developmental temperature and/or an accumulation of thermal injury (discussed below).

We found substantial differences in adult body size based on rearing temperature, wherein individuals reared at 22°C had approximately 10% larger wing lengths than those reared at 30°C. This finding aligns with the temperature-size rule—one of the most consistently observed rules in biology, positing that ectotherms reared at cooler temperatures grow more slowly but achieve larger adult body sizes (72, 73)—and has previously been observed across mosquito species (74–77). Prior work on *Ae. sierrensis* specifically has shown that wing to body length ratios are consistent across populations, suggesting our use of wing length is a valid measure of body size (49). Larger adults, in turn, are typically able to endure longer durations of thermal stress due to slower rates of resource depletion and water loss under stress, and higher thermal inertia (36, 78–81). Accordingly, prior studies in ectotherms, including *Aedes*, have found that larger adult body sizes are associated with higher upper thermal limits, as measured by longer knockdown times (49, 80, 82–84). Our results suggest a link between these prior findings, with warmer developmental temperatures leading to smaller adult mosquito body sizes, which may in turn drive the observed reduction in acute heat tolerance (though we note that wing length alone was not a significant predictor of knockdown time in our experiment).

Our finding of lower acute heat tolerance following heat exposure during rearing could also be due to an accumulation of thermal injury, whereby physiological stress incurred during prolonged heat exposure at the juvenile life stage compromises acute responses to thermal stress at the adult life stage. In our experiment, survivorship differed markedly between treatments, with approximately 18% of heat-selected individuals surviving from larvae to adulthood compared to 54% of control individuals, suggesting that high temperature exposure during rearing caused differences in survival probability (and thus fitness) in our single, genetically-diverse, source population. Sublethal high temperature exposures are well known to reduce ectotherm trait performance (85–89), and the negative impacts of thermal stress may be cumulative (90, 91). Whether high temperature exposure results in thermal acclimation versus injury depends, in part, on the intensity and duration of heat experienced (81, 87, 90–95)—a finding that has led to varying estimates of upper thermal limits in ectotherms based on experimental methodology (*e.g.*, static versus dynamic thermal knockdown assays). In our experiment, surviving individuals from the heat-selected group experienced several days at 30°C—a temperature that may be close to their upper thermal limits for prolonged larval survival (17). We transferred all surviving individuals to control temperatures (22°C) upon pupation, thereby providing several days for heat injury repair and recovery prior to the knockdown assay. However, despite surviving prolonged heat exposure, individuals may have incurred heat damage that was irreparable and/or that required substantial energetic allocation to repair, a mechanism supported by the lower survival of pupae that experienced heat treatment as larvae. Differentiating between these potential mechanisms—body size and energetic reserve variation and thermal injury—which are non-mutually exclusive, will ultimately require rearing individuals across thermal environments for several generations (*e.g*., (25)).

### Polygenic architecture of prolonged and acute heat tolerance

Our investigation of the genomic architecture of thermal tolerance used a de novo chromosome-level reference genome assembly for *Ae. sierrensis* (1.183 Gb, available at NCBI BioProject ID: PRJNA1119052 upon manuscript acceptance) and revealed a polygenic architecture for both prolonged and acute heat tolerance. That is, we identified hundreds of candidate single nucleotide polymorphisms (SNPs) distributed across the *Ae. sierrensis* genome that were associated with surviving prolonged heat exposure or resisting acute thermal stress. These candidate SNPs were identified using both genomic differentiation (F_ST_) and genome-wide association (GWA) approaches, and rigorous quality filtering to reduce false positives. Further, they displayed significantly larger differences in frequency between groups (*i.e.*, heat-selected versus control individuals or short versus long knockdown times) than did control SNPs of similar starting frequency and chromosomal position, strengthening inference of their association with thermal tolerance.

The genomic regions with an elevated signal of SNPs associated with thermal tolerance could indicate regions of selection in which structural genomic changes, such as chromosomal inversions, insertions, deletions, and/or duplications, are present. As inversions have previously been implicated in mosquito climate adaptation and pathogen infection susceptibility (37, 51, 96), we quantified their potential role in the genomic patterns of selection observed in our experiment. We found inversions to be putatively pervasive within the *Ae. sierrensis* genome, whereby we identified approximately 450 inversions between 1-200 Mb in length and at >5% frequency segregating in our focal populations. These were enriched in the specific chromosomal regions where we observed a dramatic elevation in the number of SNPs associated with larval heat tolerance (Figure 3A, Supplemental Methods). Further, five inversions within this region exhibited systematic frequency differences between control and heat-selected larvae, suggesting their role in mosquito heat stress responses. This is consistent with a large body of literature, including in *Anopheles* spp., finding that inversions are an important mechanism of ecological adaptation as suppressed recombination between the inversion breakpoints can lead to co-adapted gene complexes and/or preserved combinations of locally adapted alleles (34–42, 97, 98). In particular, the acquisition of inversions *2La* and *2Rb* through introgression from *An. arabiensis* is thought to have enabled *An. gambiae* to expand its ecological niche and become the dominant malaria vector in much of sub-Saharan Africa (38, 39, 99–101). Similarly, several chromosomal inversions in *An. funestus*—an additional key malaria vector in tropical Africa—were found to underscore adaptation to an anthropogenic larval habitat (irrigated rice fields), enabling niche diversification that may challenge vector control efforts (37). Investigating the extent to which thermal tolerance interacts directly and indirectly with vector competence is an intriguing area of future research. In particular, whether warm-adapted genotypes are more or less susceptible to becoming competent vectors—a dynamic that could alter disease transmission patterns under warming—is unknown. Finally, we note that the short-read sequencing data used in this analysis is not ideal for calling structural variants (though inversions with systematic differences between treatments are unlikely to be technical artifacts), and the use of long-read sequencing technology and/or cytogenetic analysis to resolve the inversions segregating in our focal species is an exciting area of future research.

By assigning the candidate thermal tolerance SNPs to genes, we found several that mapped to genes previously associated with responses to heat or other environmental stressors in a range of ectotherms (Supplemental Table S7). In particular, genes associated with prolonged heat tolerance in our experiment included histone H3, previously implicated in heat shock memory and enhanced survival under subsequent heat exposure (53, 54); profilin, an actin-binding protein that may function as a molecular chaperone and play an important role in the heat stress response (55, 56); and cytochrome P450, a class of proteins implicated in the thermal stress response in plants (57), corals (58), and insects (59). We also identified several genes that were associated with both prolonged and acute heat tolerance, exceeding expectations of genomic overlap due to chance (*i.e.,* 6 observed overlapping genes versus ∼2 expected out of a total of 493 candidate genes). These overlapping genes mapped to genes previously found to be involved in DNA damage repair in ectotherms (‘DNA repair endonuclease’)(102–104), and sensory perception in *Ae. aegypti* (‘dopamine receptor-1’)(105) or *Ae. albopictus* (‘sensory neuron membrane protein 2’)(106). As the candidate genes identified here largely align with prior associations of abiotic stress resistance in other ectotherms, they may be valuable targets for future transcriptomic and functional studies to clarify their role in mosquito heat tolerance.

### Potential to adapt on pace with warming

Our data suggest that natural mosquito populations may harbor the potential to adapt on pace with climate warming, and thus incorporating this adaptive potential is critical to accurately projecting the range of mosquitoes and other disease vectors under future climates. In particular, we parameterized a simple evolutionary model estimating the maximum potential rate of evolutionary change in larval thermal tolerance in comparison with expected rates of change in mean temperatures during the spring, the larval activity period. We focused on larval heat tolerance as recent evidence suggests this may be the bottleneck to thermal adaptation in this species (17). We found that, under most parameter values informed by our experimental and genomic data, estimated rates of adaptation exceeded projected rates of warming in mean spring temperatures under a moderate warming scenario (*i.e.*, RCP 4.5). This suggests that the warm edge limits of the species (and similar disease vector congeners) may not contract as quickly as assumed in most models, and the overall suitable range for the species may increase under global warming.

We note that our evolutionary model did not incorporate the presence of daily and seasonal temperature variation, concurrent stressors in other abiotic or biotic factors (*e.g.*, drought, resource availability, land use change, human insecticide applications), or phenotypic plasticity, which may alter rates of adaptive evolution (61, 107–110). In particular, phenotypic plasticity may be a key mechanism of mosquito responses to warming, particularly to short-term thermal extremes (22, 25, 48, 111), and may trade off with basal heat tolerance (112–114). However, the extent of phenotypic plasticity in natural mosquito populations and its relationship to basal thermal tolerance and adaptive potential was not directly explored here and remain poorly understood. Another limitation of this modeling approach is that the strength of selection was simply estimated as the difference in survival between our experimentally-imposed temperature treatments. These temperatures—22 and 30°C—were chosen to approximately capture the current maximum temperature this population may experience during the larval activity period and the upper thermal limits for larval survival, respectively, but are not specific to the projected shift in temperatures that natural populations may experience. That is, natural populations currently experience temperature variation on multiple time scales and may experience future warming as gradual, punctuated, and/or accelerating changes in temperature over time, each of which may impose different strengths of selection which could be lesser or greater than our estimate (115–117). Similarly, we also assume that the strength of selection, and responses to selection are uniform across space. That is, we do not consider environmental gradients in selection, potentially negative effects of gene flow on adaptive capacity (*i.e.,* “swamping”), or reduced genomic variation in range-edge versus central populations—each of which may alter, and likely reduce, the realized adaptive potential (118, 119). Lastly, our model considered only evolutionary adaptation in prolonged heat tolerance at the larval stage, which may have ecological trade-offs, and/or a distinct genetic underpinning from other heat tolerance traits. In general, evolutionary models such as that used here have rarely been validated in natural settings, thus we interpret these as results under idealized conditions that warrant further investigation under more ecologically realistic settings.

Despite these caveats, the evidence for climate adaptive potential presented here aligns with several prior studies finding high levels of phenotypic or genomic variation in heat tolerance in natural mosquito populations (24, 25, 39, 120), phenotypic shifts in heat tolerance over time (25), and rapid genomic shifts when invading novel climates (60). Collectively, these findings provide compelling evidence that evolutionary adaptation could enable mosquito populations to persist in regions where they are otherwise expected to decline due to warming. For example, projections based on fixed mosquito heat tolerance predict declines in *Aedes albopictus*-transmitted arboviruses in the tropics (5) and in *Anopheles gambiae*-transmitted malaria in western Africa (121) in coming decades. Our findings suggest that, if these species have similar levels of standing genetic variation in thermal tolerance as our focal *Ae. sierrensis*, these predictions may underestimate the habitable range of mosquito species and mosquito-borne disease transmission under climate warming.

## Materials and Methods

### Source population

Larval *Ae. sierrensis* were collected by Solano County Mosquito Abatement District personnel from various tree holes in Solano County, California in spring, 2019 (Figure 1, Supplemental Figure S11). Average daily temperatures in this region during the spring (here defined as January - April) are approximately 10 - 15°C and average daily maximum temperatures are approximately 14 - 22°C respectively (122). Collected larvae were reared for two generations at 20 - 22°C at the Solano County Mosquito Abatement District to minimize direct environmental effects on thermal tolerance (*i.e*., ‘phenotypic plasticity’) and maternal/cross-generational effects. Approximately 300 females from the F_2_ generation were blood-fed and produced eggs for use in the experiment. Prior to experimentation, eggs were transported to the Mordecai lab at Stanford University and maintained at 20 - 22°C and 6h/18h light/dark cycles to prevent diapause. Our focus on patterns of variation within a single population allowed the identification of trait variation to be minimally impacted by population substructure, which can drive spurious signals in association-based studies (123). We further selected our focal population to be from the center of the species range, and thus most likely harboring mutations present across the range and insensitive to idiosyncratic patterns of diversity that can accumulate at range edges (124).

### Selection experiment set-up

The selection experiment began with *Ae. sierrensis* at the egg stage (Figure 1). Egg paper containing approximately 200 eggs each were hatched in plastic trays containing 1L boiled distilled water cooled to room temperature and 1 tablespoon larval food (4 parts high protein cat chow: 4 parts alfalfa pellets: 1 part nutritional yeast). All eggs were hatched at 22°C under 14h/10h light/dark cycles. Upon hatching, individual larvae were then designated to either the control or treatment group, which each consisted of four identical replicates (Figure 1). Individuals were randomly assigned to a replicate such that approximately 30% of larvae were designated to replicates in the control group and 70% to replicates in the treatment (‘heat-selected’) group. This uneven assignment was due to expected reduced survivorship in the heat-selected groups, as observed during pilot experiments. Control group individuals were maintained at 22°C through to adulthood (approximately 18 days after hatching). Treatment group larvae were placed in incubators that were ramped from 25 to 30°C over the course of 3 days. This ramping period was used because prior pilot experiments in the lab found high larval mortality (>95%) when transferring 1^st^ instar larvae directly from 22 to 30°C. These specific temperature treatments—22 and 30°C were specifically chosen to approximate the currently daily maxima for this population in the spring, when larvae are developing, and the upper thermal limits for larval survival as measured under constant temperature conditions (17). To reduce accumulated thermal stress across the lifetime, treatment group larvae were transferred to the control temperature (22°C) upon pupation (approximately 14 days after hatching) and remained here through adulthood (approximately 4 days after pupation). Thus, our selection specifically aimed to target prolonged thermal tolerance at the larval stage, leading to genotype frequency differences between the control and heat-selected adults that reflect this early life-history selection event. Individuals from each replicate of the control and treatment groups were maintained in plastic cups and fed 1 teaspoon larval food every three days. Once reaching the adult life stage, individuals were transferred to breeding cages (BioQuip). The heat-tolerance assay was performed on adults that had eclosed 48-72 hours prior and had not been sugar of blood-fed. This procedure—from hatching to knockdown assay—was conducted three times to ensure our results were robust to minor variation in laboratory experimental conditions.

### Heat-tolerance assay

We used a thermal knockdown assay to assess the upper heat tolerance of adult *Ae. sierrensis* from the control and treatment groups. This is a commonly used assay to measure upper thermal limits of arthropods, including mosquitoes, and has been shown to accurately predict insect species distributions in the field and to be a relevant proxy for fitness under heat stress (17, 24, 25, 47, 48, 125), (but note that methodological details such as the initial and final temperature conditions have varied between studies). We followed the thermal knockdown protocols of Mitchell *et al.* 2011 (126) and van Heerwaarden *et al.* 2016 (127). Specifically, adult *Ae. sierrensis* were placed in individual 5-mL plastic vials and immersed in a water bath set initially to 28°C. After a 15-minute acclimation period, the water bath temperature was increased to 38°C at a rate of 0.5°C per minute (128). Heat tolerance was scored as the ‘knockdown time’—the time after immersion at which an individual loses muscle function and can no longer right itself from a dorsal position. This assay thus represents the acute thermal tolerance of all control individuals and individuals that had survived prolonged exposure to thermal stress at the larval stage. Immediately after knockdown, samples were placed in individual tubes and stored at −80°C. Thermal knockdown assays were performed on adults that had eclosed 48 - 72 hours prior and had not been sugar or blood fed. Assays were performed on eight adults at a time and the same observer performed all assays.

### Body size estimates

To estimate the body size of each individual used in the experiment, we measured mosquito wing length—a commonly used proxy, including in *Aedes* spp. mosquitoes (129–134). The wing length-body size relationship has been validated in *Ae. sierrensis* specifically, and corroborated with measurements of thorax lengths, with no systematic differences identified in wing to thorax length ratios across temperatures (49). Prior to DNA extraction, we detached the left wing using fine forceps and attached it to a microscope calibration slide. We then took an image of the microscope field and used *ImageJ* (135) to measure the length of the wing as the distance from the alula to the wing tip, excluding the wing scales (Supplemental Figure S8)(136, 137). We repeated each measurement and herein report the average of these two measurements. We excluded any wings that were damaged during removal or the knockdown assay.

### Statistical analysis on knockdown assay

To investigate variation in knockdown times by treatment, we used a linear mixed-effects model implemented using the ‘lme4’ package in R (138). We used knockdown time as the outcome variable and treatment and sex as categorical fixed effects. To account for potential variation between biological replicates (as the experiment was repeated in three rounds) and knockdown assays, we included these as random intercepts in the model.

### DNA extraction and sequencing

We extracted DNA from each individual using an AllPrep DNA/RNA kit (Qiagen, Valencia, CA), according to the manufacturer’s protocol. We quantified the extracted DNA using a Qubit and assessed the quality using Nanodrop. As the Nanodrop results indicated low 260/230 ratios, we performed a 1.8x clean-up using Illumina Purification Beads. Following quantification, we proceeded with library preparation using approximately 25ng genomic DNA per sample. We prepared libraries using the Illumina DNA prep kit, following manufacturer’s protocols. The libraries were pooled and sequenced as 150-bp paired-end reads on four lanes (58-59 samples / lane) of a NovaSeq 600 Illumina at the Stanford Genome Sequencing Service Center. Summary statistics for each sample are available in Supplemental Tables S4-S5.

### Reference genome assembly

We assembled a de novo reference genome for *Aedes sierrensis* to facilitate genomic analysis in the absence of a previously available reference for this species (available at NCBI BioProject ID: PRJNA1119052 upon manuscript acceptance)(Supplemental Methods). Briefly, we selected a single adult female *Ae. sierrensis* that was field-collected from Eugene, Oregon for PacBio HiFi sequencing. After high molecular weight genomic DNA was extracted, two libraries were prepared from this sample—a Low and Ultra-Low DNA library—each using the SMRTbell Express Prep kit and following manufacturer’s protocols. Each library was loaded onto a separate 8M SMRT Cell and sequenced on a Sequel II System at the University of Oregon Genomics and Cell Characterization Core Facility.

We then assembled the genome using default parameters in Hifiasm v 0.16—a haplotype resolved assembler optimized for PacBio HiFi reads (Cheng *et al.* 2021). Evaluating the assembly for missing or duplicated genes indicated a high level of completeness (97.1%) and duplication rates on par with that of recent de novo assemblies in other mosquito species (Supplemental Methods)(139–141). We used the *Aedes aegypti* Aaeg L5 genome (NCBI BioProject ID: PRJNA940745) to scaffold the draft assembly into chromosomes using RagTag (142), and found that 96% (1.139 Gb) of our assembly scaffolded to this Aaeg L5 genome.

### Genome annotation

We first identified and masked repetitive elements in our reference genome assembly using RepeatModeler v2.0.1 with a custom repeat library (143) and RepeatMasker 4.1.6. (144). We then annotated the genome for protein-coding genes using *BRAKER2*—a fully automated pipeline that uses the tools *GENEMARK-ES/ET* (145) and *AUGUSTUS* (146) for gene structure prediction (147). Specifically, we conducted *ab initio* gene prediction in BRAKER2 v 2.1.6, using the genome file only (*i.e*., without additional evidence from RNA-Seq or protein data, as these are unavailable) and a minimum contig length of 10,000.

### Read trimming and variant calling

Raw reads were first quality filtered and trimmed using *Trimmomatic* V0.39 (148) with the following parameters: ILLUMINACLIP:TruSeq3-PE.fa:2:30:10 LEADING:3 TRAILING:3 MINLEN:35 SLIDINGWINDOW:4:15. We then aligned these reads to the scaffolded reference genome using *BWA-MEM* v0.7.12, with default parameters (149). We marked and removed duplicate reads using *picard* v2.0.1. We then identified single nucleotide polymorphisms (SNPs) in our samples using *bcftools* v1.18 (150) and filtered variants using *vcftools* v0.1.16 (151) with the following parameters: minor allele frequency of 0.05, minimum depth of 10x, minimum average quality of 40, and a maximum variant missing of 0.995. We then filtered out any SNPs with multiple alleles using *bcftools*, keeping only bi-allelic polymorphisms. This retained 3,564,483 SNPs. As large linkage blocks have been found in related *Aedes* species (152), we filtered for linkage disequilibrium (LD) in this SNP set using an LD-based SNP pruning algorithm in *plink* v1.9 (153). Specifically, we used a sliding window of 50 SNPs, a window shift increment of 5 SNPs, and variance inflation factor (VIF) of 1.5, which corresponds to an R^2^ of 0.3 for the focal SNP regressed against all other SNPs in the window (154, 155). This yielded 583,889 independent SNPs that were retained for downstream analysis.

### Population diversity metrics

To estimate the genetic diversity of our starting population, we calculated the individual-level heterozygosity and population-level nucleotide diversity (Π) in 10 kb windows using *vcftools* v0.1.16 (151). We estimated these metrics using only the control individuals that survived to adulthood, as representative of the population prior to heat selection.

### Identifying genetic variants associated with heat tolerance

To investigate the genetic basis of heat tolerance, we used a combination of principal components analysis (PCA)(Supplemental Figures S9-S10), statistical tests of genomic differentiation (F_ST_), and genome-wide association (GWA) approaches. We first visualized overall genomic variation through PCA on the allele frequency matrix (after centering and scaling) using the *prcomp* function in R. We used these PCA visualizations to briefly explore whether there was a dominant signal of treatment or sex, as well as experimental round, as a quality control measure (Supplemental Figures S9-S10). We then detected candidate SNPs underlying heat tolerance using the following approaches: 1) genomic differentiation (F_ST_) between control and heat-selected individuals, 2) a case-control GWA analysis between control and heat-selected individuals, and 3) GWA using knockdown time as the phenotype. Approaches one and two are aimed at identifying genetic variants associated with tolerance to prolonged heat exposure during development. We adopted two independent but complementary approaches in order to compare SNPs identified under the varying assumptions and approaches of each method and to ultimately refine the candidate SNP list (*i.e*., SNPs identified by both approaches are less likely to be false positives). Approach three is designed to detect variants associated with acute heat tolerance at the adult life stage using a standard GWA approach (*i.e.*, regression of continuous trait value on genotype status).

Approach one (F_ST_-based approach), was implemented using R package *OutFLANK* v0.2, which calculates F_ST_ at each SNP using the Weir and Cockerham method (156) then identifies SNPs that deviate from an inferred neutral F_st_ distribution (157). We considered SNPs below a q-value threshold of 0.05 and with F_st_ > 0.05 as candidate SNPs putatively underlying heat tolerance. Our second approach to identify SNPs associated with prolonged, larval thermal tolerance was a modified GWA, whereby the ‘phenotype’ used in the regression was the treatment of the sequenced individual. We implemented a logistic regression and included sex as a covariate to control for sex-specific variation in mosquito heat tolerance (Andersen *et al.* 2006), using *plink* v1.9 (153). Finally, we conducted a standard GWA for adult, acute thermal tolerance using knockdown time as the phenotype, including sex as a covariate as above, and additionally including heat selection treatment and wing length as covariates to account for effects of larval rearing temperatures and body size on acute adult heat tolerance (71)(also conducted using *plink* v1.9). To correct for residual linkage disequilibrium in both GWA-based approaches, we performed an LD-based ‘clumping’ procedure, wherein SNP-based results from the association analyses are grouped based on estimates of LD between SNPs. We implemented this procedure in *plink* using default parameters (*i.e.*, 0.0001 significance threshold for the focal SNP, 0.01 significance threshold for clumped SNPs, 0.50 R2 threshold for clumping, and a 250 kb window for clumping). We defined candidate SNPs as those with p < 0.01 after Benjamini-Hochberg false discovery rate (FDR) correction.

To identify the genes associated with these candidate SNPs, we assigned SNPs to genes based on their position and the BRAKER gene annotation. For SNPs that did not fall within a BRAKER-annotated gene, we assigned it to the closest gene if this was within 50 kb, otherwise we removed it from downstream analysis on candidate gene overlap and function.

Using the candidate gene list from each approach, we then investigated the genomic basis of prolonged versus acute heat exposure. Specifically, we compared the genes identified through F_ST_ or GWA on control and heat-selected individuals (*i.e.,* representing prolonged heat exposure), and through GWA on knockdown time (*i.e.,* representing acute heat exposure). Next, we sought to determine whether the number of shared genes identified by these approaches was more or less than that expected by chance, to identify candidate genes identified by multiple independent approaches, and determine whether or not the pathways related to heat tolerance between life-stages (and at long-term vs acute scales) were similar. To do so, we developed a null distribution of gene overlap by drawing random samples from the available gene set wherein, for each focal gene, we selected a random gene that was a) on the same chromosome and b) within one standard deviation of gene length. We did this for the focal genes identified through each approach, then determined the number of overlapping genes in each of 500 iterations to generate a null distribution of gene overlap. If the true overlap between genes identified in each approach was greater than the expectation based on this null distribution, we inferred that the approaches were identifying shared pathways.

Lastly, to identify the putative biological function of focal genes identified herein, we mapped the gene sequences to annotated transcriptomes of related *Aedes* species (*i.e*., *Ae. albopictus,* NCBI accession: GCF_006496715.1; *Ae.aegypti,* NCBI accession: GCF_002204515.2) using BLASTN. In the case of multiple hits for a given sequence, we used the result with the lowest E-value and highest Max score.

### Investigating structural variation

We used four short-read structural variant callers, *Manta* v1.6.0, *Delly2* v1.2.6, *Lumpy* (via *smoove* v0.2.8), and *GRIDSS2* v2.13.2, to identify inversions in each of our samples (158–161). *GRIDSS2* was run with the flag --skipsoftcliprealignment, while the other three callers were run with default settings. For each sample, the sets of variants from each of the four callers were merged together with *Jasmine* v1.1.5, requiring support from at least two callers to keep a variant in the final set for that sample (using the min_support=2 argument)(162). Finally, variant sets from all samples were merged into a single, population-level VCF with *Jasmine*. In both instances, *Jasmine* was run with the arguments spec_reads=8, spec_len=35, --dup-to-ins --mark-specific, and --normalize-type. To minimize spurious detection (*e.g.*, due to misalignment of transposable elements or repetitive sequences), the full set of variants across samples was filtered to only include inversions of size 1 to 200 Mb and frequency >5% in the population (n=444) with *bcftools* v1.17 and custom bash scripts.

To identify inversions that may be associated with heat tolerance, we compared the frequency of each inversion between either the control and heat-selected individuals or between individuals from the top 25% and bottom 25% of knockdown times (controlling for treatment and sex) using a Chi-squared test. We then compared the observed Chi-squared test statistic to that obtained when comparing inversion frequency after randomly shuffling the group labels (n = 500 permutations). We considered inversions as significantly differentiated between groups if their observed Chi-squared statistic was significantly greater than expected from the permutations at p<0.05 after Bonferroni adjustment for multiple testing. Lastly, to compare the genomic position of inversions and candidate SNPs, we investigated whether inversions occurred in specific chromosomal regions in which we observed an elevated number of SNPs associated with larval heat tolerance (Figure 3A, Supplemental Figure S3). These regions of interest were explicitly defined based on SNP signals within sliding windows following methods in Rudman et al. 2022 (163)(Supplemental Methods).

### Estimating allele frequency shifts

For candidate loci underlying differences between control and heat-selected individuals, we investigated shifts in allele frequencies between these groups, relative to a set of matched controls. For each focal SNP, we generated a random set of 10 matched control SNPs with the following criteria: presence on the same chromosome, +/− 2.5% baseline allele frequency (*i.e*., allele frequency in the control individuals), and at least 100 kb away from the focal SNP (to account for linkage disequilibrium). We then compared differences in the distribution of allele frequency shifts in the focal SNPs relative to their matched controls using a Kolmogorov-Smirnov (K-S) test, and compared shifts in allele frequency based on starting median allele frequency (MAF)(Supplemental Figure S4). We repeated this process to investigate allele frequency differences between individuals with high (upper 50% of phenotypic distribution) or low (bottom 50%) knockdown times relative to their treatment and sex. Herein, for each focal SNP identified from the GWA on knockdown time, we generated a set of 10 matched controls based on presence on the same chromosome and at least 100 kbps separation from the focal SNP.

### Estimating adaptive potential

To investigate whether the standing variation in thermal tolerance observed here may enable adaptation to climate warming, we used an evolutionary rescue model framework (15, 61, 62). These models compare the maximum potential rate of evolutionary change for a population to the projected rate of environmental change. If evolutionary rates exceed that of environmental change, populations may persist through evolutionary adaptation (110, 164–168). Evolutionary rescue models have provided useful estimates of climate adaptive potential across a variety of taxa (169–172). However, there are few examples of their validation in natural settings, thus we pose this analysis as a means of estimating adaptive potential under idealized and simplified conditions that warrant further investigation under more ecologically realistic settings, rather than an attempt to estimate a precise rate of warming to which mosquito populations may adapt. Here, we specifically consider the rate of evolutionary change in the thermal tolerance of larval survival and compare it to rates of change in mean daily temperatures in the larval activity period (January - April) as projected under a moderate warming scenario for the southern portion of the *Ae. sierrensis* distribution (discussed further below). We focus on larval thermal tolerance as our recent investigation of thermal tolerance across the life stages of this species indicate that larval survival may be the bottleneck to adaptation (17). We use the analytic, quantitative-genetic formulation of the evolutionary rescue model below, based on Lynch and Lande 1993 and Chevin *et al.* 2010 (61, 62)(see Supplemental Methods for derivation).

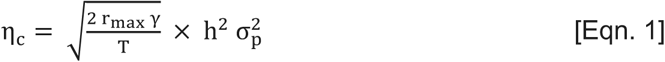

Here, η_c_ is the maximum rate of environmental change under which the population could persist (which is equivalent to the maximum rate of adaptive evolution), r_max_ is the maximum rate of population growth under optimal conditions, γ is the strength of selection, T is the generation time, h^2^ is the heritability of the trait, and 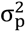 is the phenotypic variance. We note that this formulation does not incorporate phenotypic plasticity, which could modify the strength of selection and rate of change in the trait under warming (61).

We parameterized the model using estimates from our experimental and genomic results, and prior estimates of maximum mosquito population growth rate and *Ae. sierrensis* generation time (Supplemental Table S8). Namely, prior studies have estimated r_max_ for *Ae. aegypti, An. gambiae*, and *Cx. pipiens* as 0.24 - 0.335, 0.187, 0.379, respectively, based on laboratory experiments that varied larval competition and/or temperature (15, 173, 174). As r_max_ for mosquito populations in natural settings remains largely unknown, we estimate adaptive potential over a range of r_max_ from 0.15 - 0.35, based on these prior estimates. We estimated the strength of selection, γ, as the difference in larval survival between treatments (*i.e*., γ = 1 - (survival in heat-selected group / survival in control group))(175) across all experimental rounds collectively (γ = 0.578), as well as each individual experimental round (γ = 0.463, 0.606, 0.590). An important limitation in our parameterization of γ is that it was not estimated under the same temperature conditions as expected under warming. That is, we estimated selection strength by comparing larval survival at 22 and 30°C, which approximately capture the maximum temperature this population may experience in the spring and the upper thermal limits for larval survival as measured under constant temperatures (17), indicating the selective regime we imposed is likely to be biologically realistic. However, this temperature differential may be larger than that experienced by natural populations in coming decades, causing us to overestimate selection strength (though this estimate may also be an underestimate during extreme heat events when temperatures far exceed expected mean shifts). However, in the absence of theoretical or experimental approaches to estimate γ under environmental conditions that are changing continuously and non-linearly with respect to time, we use the estimate of γ made here and interpret our results cautiously. To estimate heritability and phenotypic variance, we used *GCTA*—a tool developed to estimate these parameters for complex traits based on genome-wide SNPs (176). First, the pairwise genetic relatedness between all individuals is estimated based on all SNPs. As the *GCTA* method relies heavily on linkage disequilibrium between SNPs, and can overestimate heritability under certain LD scenarios, we performed this step using the SNP list both before and after LD-pruning (see Methods: *Read trimming and variant calling*). The resulting genetic relationship matrix is then used to estimate the variance in the larval heat tolerance phenotype explained by the SNPs using restricted maximum likelihood. Herein, all SNPs are used, rather than solely those identified as focal SNPs from F_st_ or GWA approaches, to avoid overestimating effect sizes (*i.e.*, the ‘Winner’s Curse’ issue in genetic association studies)(163). We included sex as a covariate in this estimation to control for any sex-specific differences in survival.

We compared the estimated maximum rate of evolutionary change based on our parameter estimates (Supplemental Table S8), to rates of change in mean daily temperature in the spring (here, January - April)—the period when larvae are developing. Specifically, we estimated the rate of warming in spring temperatures between 2020 and 2050 under a moderate warming scenario (Representative Concentration Pathway (RCP) 4.5) across the southern portion of the *Ae. sierrensis* distribution, based on California vector surveillance data for the past decade (Supplemental Figure S11). Surveillance data were obtained from the CalSurv Gateway through data request 000045 approved on October 19, 2020 by the California Vectorborne Diseases Surveillance System. We consider this to be the most ecologically relevant metric of warming, but to ensure our results were not specific to this precise metric of warming, we also considered alternative metrics based on maximum daily spring temperatures and mean annual temperatures across the southern portion of the *Ae. sierrensis* distribution as projected under RCP 4.5, annual mean temperature across the southeastern U.S. under RCP 4.5 and RCP 8.5 (178), and recently observed rates of warming in annual mean temperatures across North America (122). For estimated rates of warming, we used temperature data from CHELSA, available at a 30 m resolution (179) or the California Basin Characterization Model, available at a 270 m resolution.

## Acknowledgments

We gratefully acknowledge Bret Barner from Solano County Mosquito Abatement District, Kristen Holt from Marina/Sonoma Mosquito and Vector Control, and Angie Nakano from San Mateo County Mosquito and Vector Control District for assisting with larval sampling and optimizing mosquito rearing protocols. We thank Kelsey Lyberger and Johannah Farner for assisting with mosquito rearing in the Mordecai lab as well as Dmitri Petrov, Molly Schumer, and members of the Moi lab for providing insights that helped shape the analytical approaches herein. This work was supported by the Philippe Cohen Graduate Fellowship, the Pacific Southwest Center of Excellence in Vector-Borne Diseases Training Grant, the Stanford Center for Computational, Evolutionary and Human Genomic Sequencing, and the NSF graduate research fellowship awarded to TOD. EAM was supported by NSF grant DEB-2011147 with Fogarty International Center, NIH grants R35GM133439, R01AI168097, and R01AI102918, and seed grants from the Stanford Doerr School of Sustainability, Woods Institute for the Environment, King Center on Global Development, and Center for Innovation in Global Health.

## Author contributions

L.I.C, E.A.M., and M.C.B. designed research; L.I.C. conducted experiments, L.I.C, T.O.D, J.A.H., B.Y.K, and M.C.B. conducted analyses, L.I.C., M.E.A., R.B.B., E.A.M., and M.C.B. interpreted data and findings; L.I.C., M.C.B, and E.A.M. wrote the manuscript. All authors reviewed and revised the manuscript.

## Competing Interest Statement

authors declare no competing interest.

## Data availability

All R scripts, bioinformatic scripts, and intermediate and final analysis files are available in an external data repository hosted on GitHub: https://github.com/lcouper/MosquitoThermalSelection

## Notes

https://github.com/lcouper/MosquitoThermalSelection

### Competing Interest Statement

The authors have declared no competing interest.

### Summary of Updates

This version includes: expanded discussion around the choice of thermal selection regime and alternative metrics of warming for comparison to restated rates of evolutionary adaptation

