## Supplemental Information for "Evolutionary adaptation under climate change: *Aedes* sp. demonstrates potential to adapt to warming"

**Affiliations:**
1. Stanford University, Department of Biology
2. University of California, Berkeley, Division of Environmental Health Sciences
3. University of California, Berkeley, Department of Integrative Biology
4. Howard Hughes Medical Institute
5. University of California, Berkeley, Department of Plant & Microbial Biology

**Corresponding author:** Lisa Couper

**This PDF file includes:**

Supporting text
Figures S1 to S11
Tables S1 to S9
SI references

**Supporting Text**

*Supporting Methods: Reference genome assembly and annotation*

To generate a reference genome, we performed PacBio HiFi sequencing on a single adult female *Ae. sierrensis* that was field-collected from Eugene, Oregon. We assembled the genome using *Hifiasm*—a haplotype resolved assembler optimized for PacBio HiFi reads (1). We first removed any remaining contaminant PacBio adaptor sequences using HiFiAdaptFilt (2). We then used *HiFiasm* v0.16 for assembly with both the low and ultra low input HiFi reads as inputs. We used the default *HiFiasm* parameters, which includes a haplotig purging step. To evaluate the assembly for missing or duplicated genes, we searched for conserved, single copy genes using *BUSCO* v5.3.0 (3) with the 'diptera_odb10' gene set—the most specific and related lineage dataset for *Ae. sierrensis*. The *BUSCO* output indicated a high level of completeness (97.1%) and duplication rates on par with that of recent de novo assemblies in other mosquito species (4–6) (Full BUSCO output: C:97.1% [S:88.5%, D:8.6%], F:0.9%, M:2.0%, n:3285). The final curated assembly consisted of 1,198 contigs of total length 1.183 Gb. Over 99% (99.52%) of the genome was contained in scaffolds > 50 Kb and the contig N50 was 113 MB.

To scaffold the draft assembly into chromosomes, we used *RagTag*—a toolset for automating scaffolding and generating chromosome-scale reference genomes (7). We used the *Aedes aegypti* Aaeg L5 genome for scaffolding (NCBI RefSeq Assembly GCF_002204515.2). Over 96% (1.139 Gb) of our *Ae. sierrensis* assembly scaffolded to the Aaeg L5 genome.

We then masked the genome for repetitive regions using *RepeatModeler* v2.0.1 (8) and *RepeatMasker* v4.1.6. (9), and annotated the genome for protein-coding genes using *BRAKER2* (10). *BRAKER2* identified 30,554 genes across the three *Ae. sierrensis* chromosomes.

*Supporting Methods: Estimating adaptive potential*

We estimated the potential for mosquito populations to adapt to warming using an analytic, quantitative-genetic evolutionary model. In particular, we use the foundational model of Lynch & Lande 1993 (11):

$$\eta_c = \sigma_g^2 \times \sqrt{\frac{2\, r_{max}}{\sigma_w^2}}$$

Herein, $\eta_c$ is the maximum rate of environmental change under which the population could persist, $r_{max}$ is the maximum rate of population growth under optimal conditions, $\sigma_w^2$ is inversely proportional to the strength of selection ($\gamma$)(12), and $\sigma_g^2$ is the genetic variance (equal to the product of trait heritability ($h^2$) and phenotypic variance ($\sigma_p^2$), by definition). Given this, we use the following, equivalent formulation in our analysis:

$$\eta_c = h^2\, \sigma_p^2 \times \sqrt{\frac{2\, r_{max}\, \gamma}{T}}$$

Herein, T refers to generation time, enabling $\eta_c$ to be expressed in units of time (*i.e.*, °C per year) instead of per generation (13). However, as the *Ae. sierrensis* generation time is approximately one year (14), the estimated per-generation and per-year values of evolutionary change are equal.

*Supporting Methods: Identifying regions with elevated SNP signal for prolonged heat tolerance*

We quantified the extent to which particular regions of the genome were enriched in SNPs driving differentiation between heat selection treatments. Specifically, using a modified clustering algorithm developed by Rudman et al. 2022 (15), we calculated significance scores for overlapping windows of 250 SNPs, with a 50 SNP step size. SNPs within each window were assigned a significance level based on the observed *OutFLANK q* and $F_{ST}$ values: significance level = 0: *q*-value < 0.2; significance level =1: *q*-value < 0.2; significance level = 2: *q*-value < 0.1 and $F_{ST}$ > 0.015; significance level = 3: *q*-value < 0.05 and $F_{ST}$ > 0.015. The score of each window was computed as the sum of significance levels for all SNPs within the window. We compared the observed scores to an empirical null distribution derived via random shuffling of SNP identities using the observed significance level values. We considered a window significantly enriched given an empirical FDR < 0.05. Overlapping enriched windows were then merged to obtain a final set of four, non-overlapping, regions enriched in SNPs differentiating heat selection treatments (one on chromosome one, one on chromosome two, and two on chromosome three). These regions of interest likely represent distinct targets of selection imposed between treatments and, due to their size (1-200 Mb), likely indicate that larger structural variants may ultimately be driving patterns of differentiation between treatments. We thus used these regions to specifically query the presence of putative inversions segregating in our study population and differentiating treatment groups (see Methods: *Investigating structural variation* and Results: *Genomic architecture of prolonged and acute heat tolerance*).

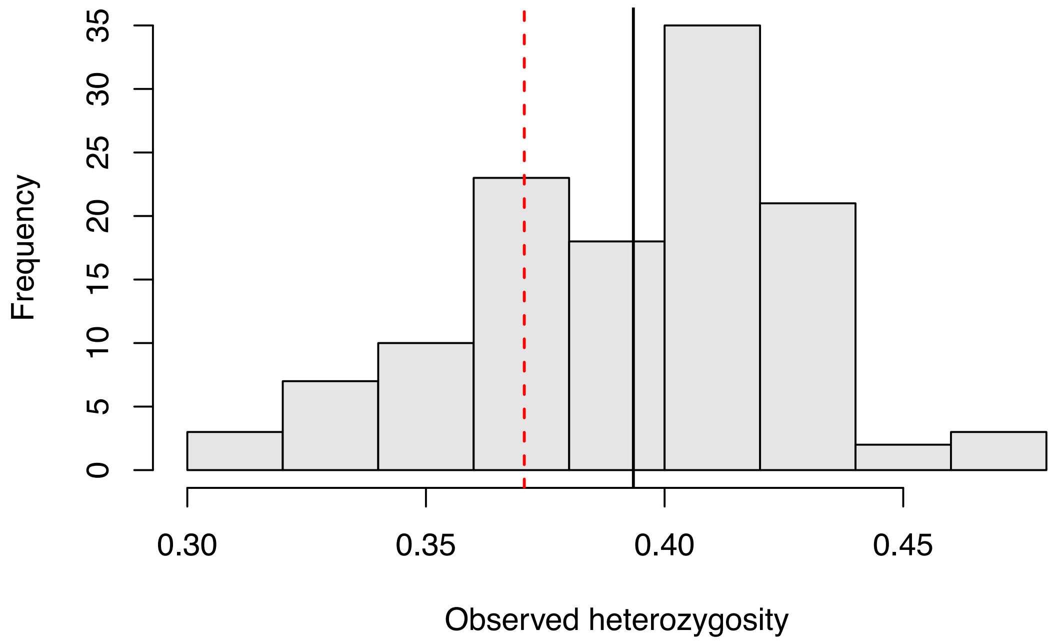

**Fig. S1.** Histogram of observed heterozygosity for all controls individuals used in the experiment. The red dashed line denotes the mean value of the expected heterozygosity (0.371) and the black line denotes the mean value of the observed heterozygosity (0.393).

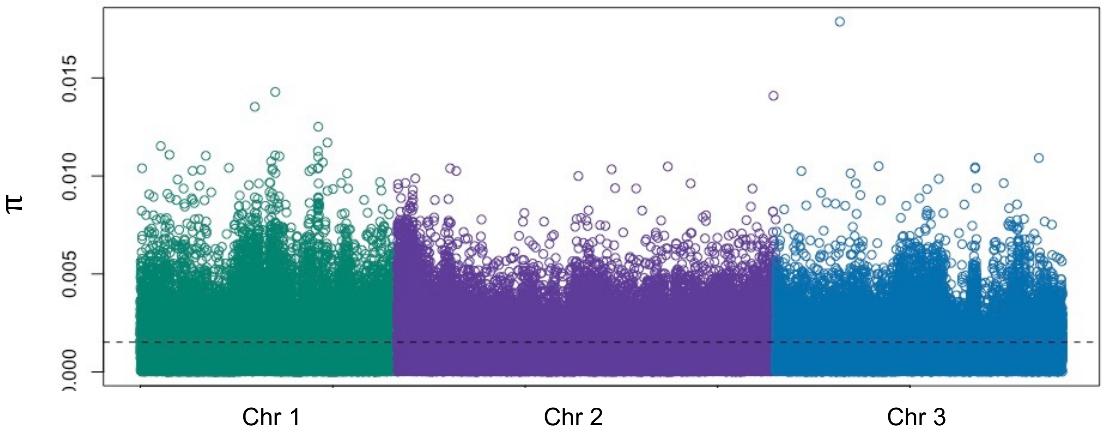

**Fig S2.** Nucleotide diversity (**π**) estimated in 10 kb windows across the genome for all individuals used in the analysis. The dashed line denotes the mean **π** (0.0015).

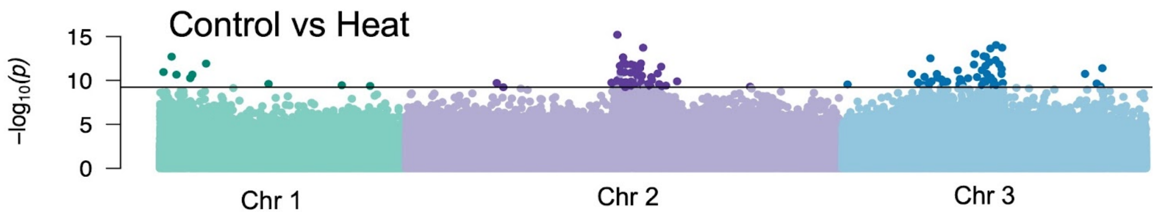

**Fig S3.** Genomic position of candidate SNPs identified through the case-control GWA between control and heat-selected individuals. The horizontal line indicates the threshold for significance as candidate SNPs (*i.e.*, FDR-corrected $p < 0.001$). Candidate SNPs are depicted as darker points in each plot. The corresponding plots showing SNPs associated with treatment based on $F_{ST}$ differentiation, or with knockdown time based on GWA are shown in Figure 3.

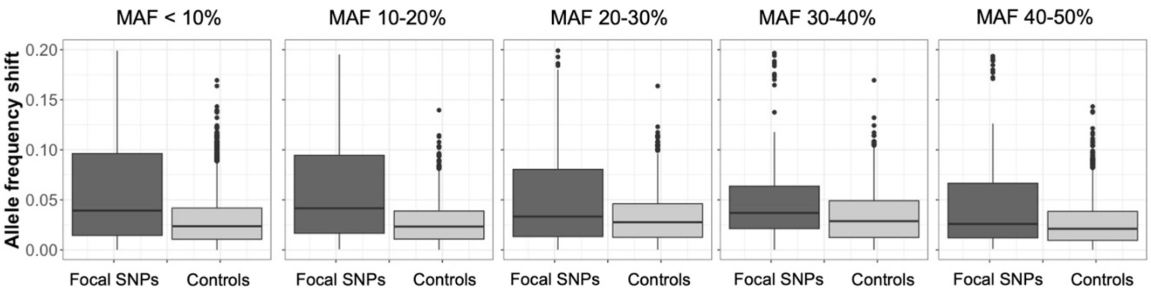

**Fig S4.** Difference in allele frequency distributions for focal SNPs (dark gray) versus their matched controls (light gray) based on the minor allele frequency in the control. Here, the focal SNPs from the $F_{ST}$ and case-control GWA approaches are shown together. The black line in each boxplot denotes the median allele frequency difference.

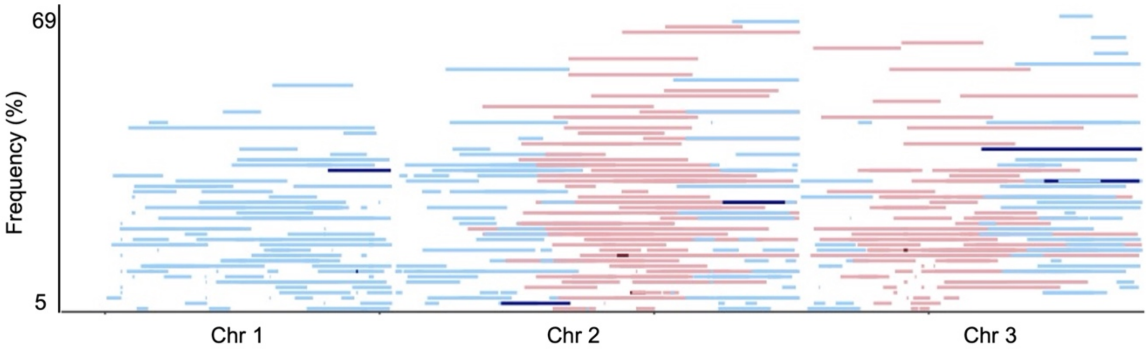

**Fig S5.** Genomic position of the 444 chromosomal inversions segregating in our focal populations (after filtering based on size between 1-200 Mb and frequency >5%). Red colors denote inversions within the specific regions in which there was an elevated number of SNPs associated with prolonged, larval heat tolerance. Blue colors denote inversions outside of this region. Darker red and blue lines denote inversions that were significantly differentiated in frequency between control and heat-selected larvae. Inversions are separated along the y-axis based on their frequency in the population.

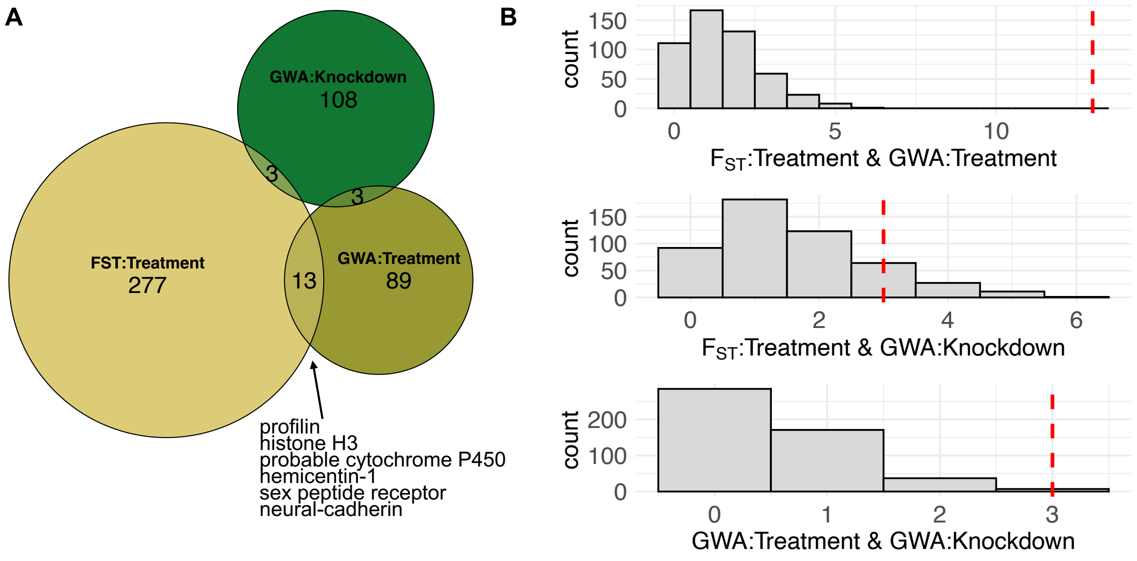

**Fig S6.** (A) Venn Diagram depicting the extent of overlap in focal genes identified through three approaches capturing prolonged ($F_{ST}$:Treatment, GWA:Treatment) and acute (GWA:Knockdown) heat tolerance traits. The names of candidate genes are listed for instances where genes have been previously characterized. (B) Histogram depicting the expected (gray bars) and observed (red dashed lines) number of overlapping focal genes from each pair of approaches.

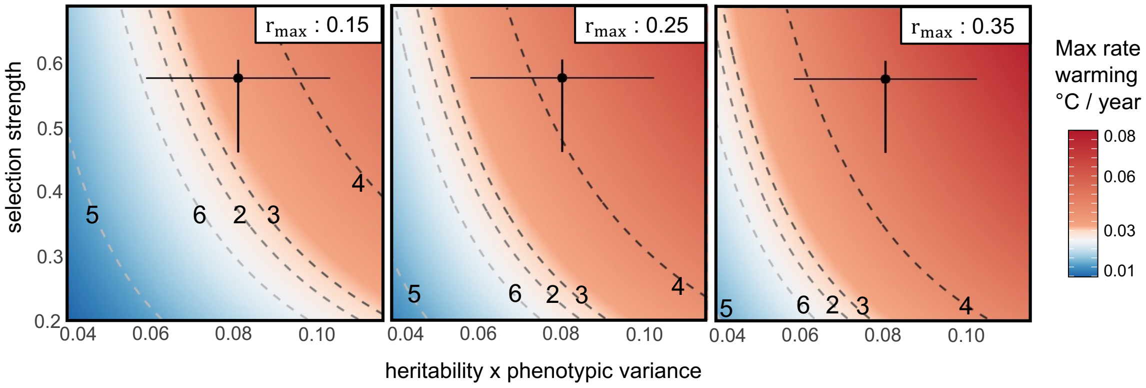

**Fig S7.** Rates of evolutionary adaptation in comparison with alternative metrics of warming. Colors denote the maximum rate of warming (°C/year) to which populations could adapt (equivalent to the model-estimated maximum rate of evolutionary change). The x-axis denotes potential values for the product of heritability ($h^2$) and phenotypic variance ($\sigma_p^2$), and the y-axis denotes potential values for selection strength ($\gamma$). The black circle on each plot denotes the point estimate for these three parameter values from our experimental and genomic data and the error bars capture the range of these parameters made under varying model assumptions (see Methods: *Estimating Adaptive Potential*). Isolines denote rates of warming based on: maximum daily spring temperatures (4) and mean annual temperatures (5) across the southern portion of the *Ae. sierrensis* distribution as projected under RCP 4.5 (0.040, 0.015 °C/year, respectively); annual mean temperature across the southeastern U.S. under RCP 4.5 (2) and RCP 8.5 (3) (0.031, 0.035 °C/year, respectively), and recently observed rates of warming in annual mean temperatures across North America (6)(0.027 °C/year). Error bars and point estimates to the right of a given isoline reflect scenarios under which the population's estimated maximum rate of evolutionary change exceeds the rate of warming. The three panels span previously estimated rates of maximum mosquito population growth rates ($r_{max}$ = 0.15, 0.25, and 0.35 for the left, center, and right panels, respectively).

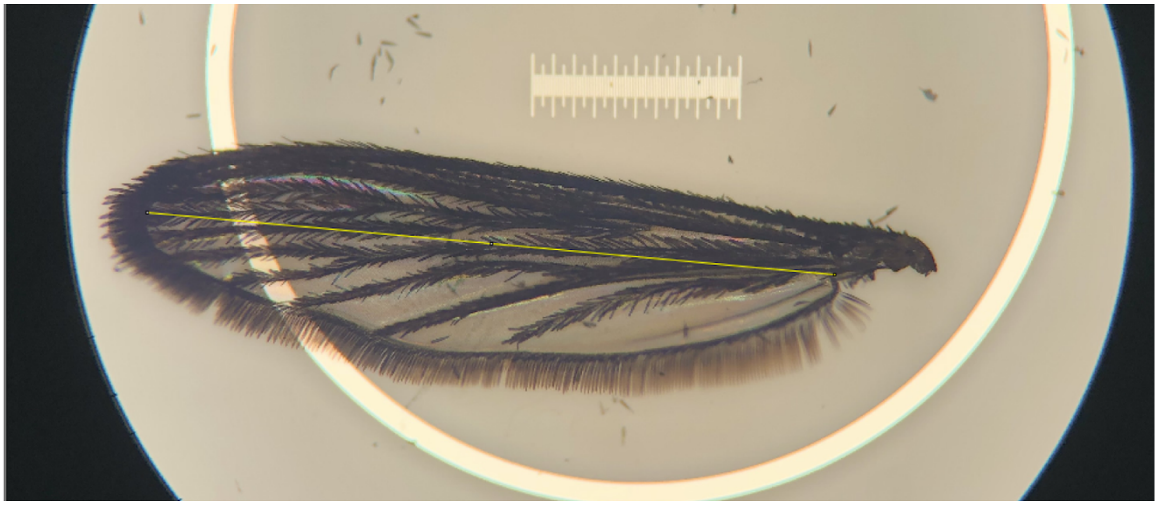

**Fig S8.** Mosquito wing length measurements. Wings were measured using ImageJ as the length from the alula to the wing tip, excluding the wing scales (indicated by the yellow line).

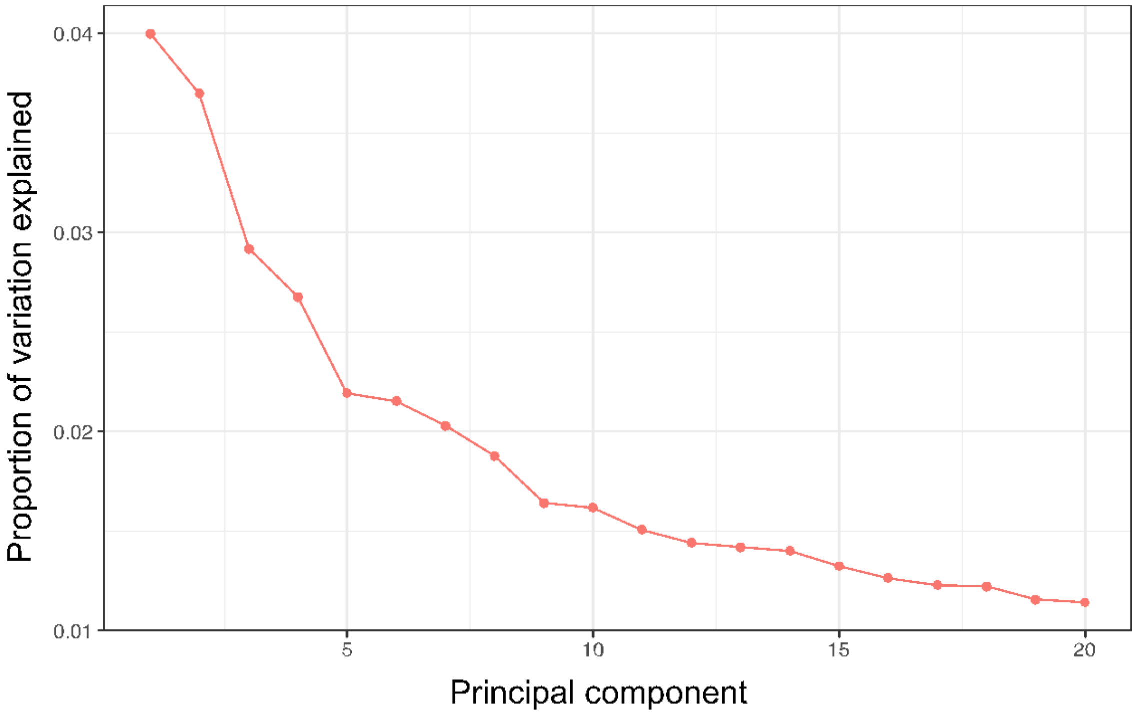

**Fig S9.** Scree plot displaying the decay in the proportion of genomic variation between control and heat-selected individuals in the first 20 principal components.

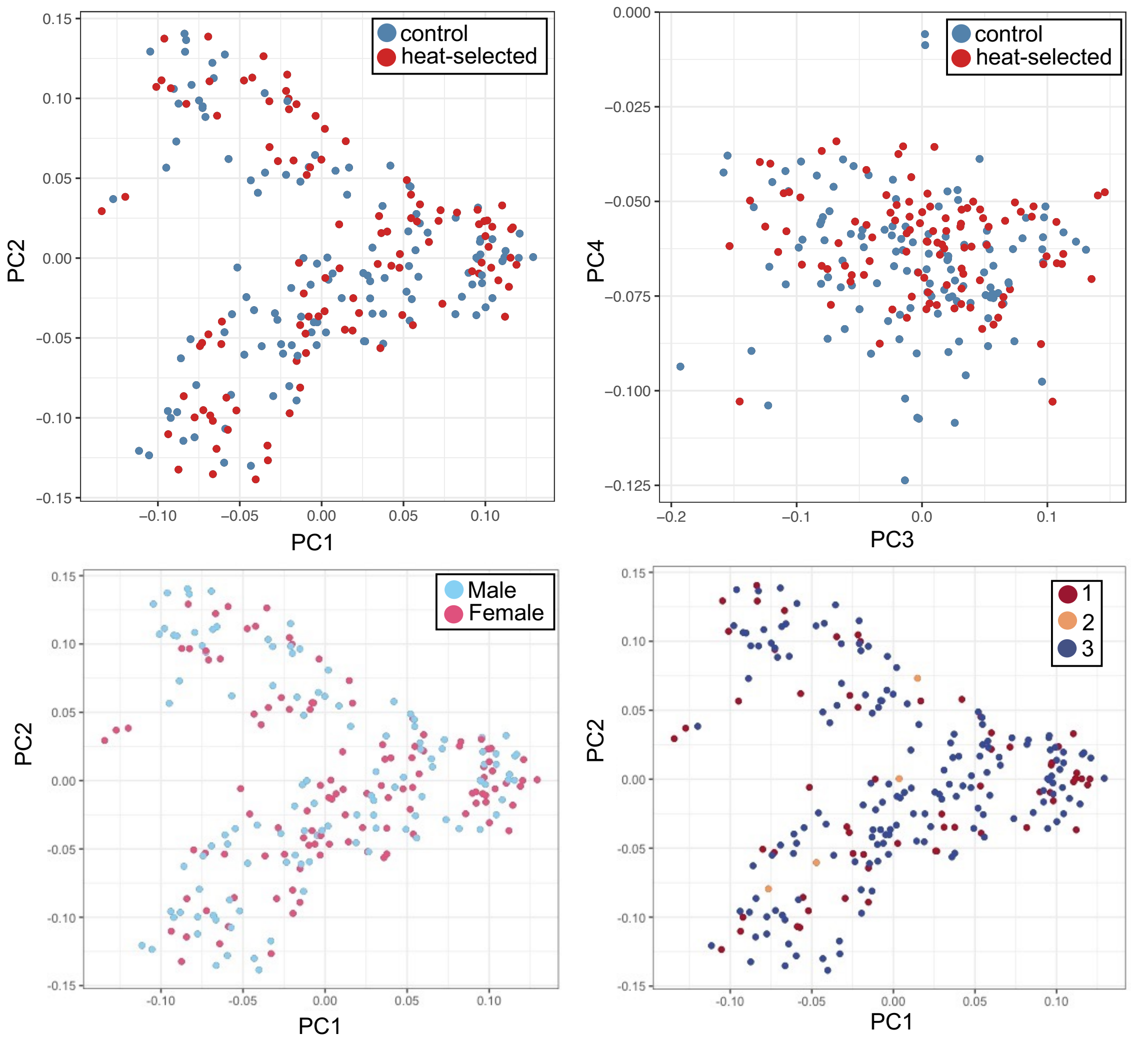

**Fig. S10.** Results of principal component analysis whereby all experimental samples are projected upon the first two principal components (top left and bottom panels) or third and fourth principal components (top right). Samples are colored based on selection treatment (top panels), sex (bottom left), or experimental round (bottom right).

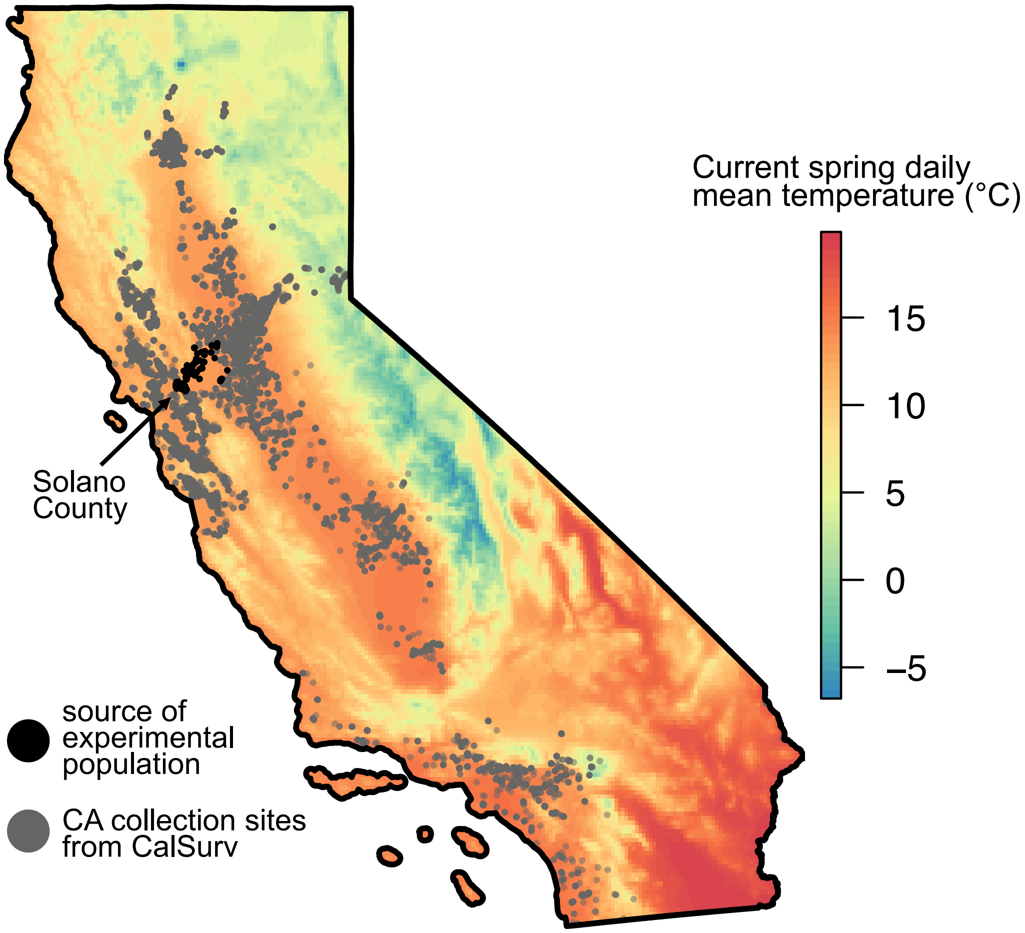

**Fig S11.** California map showing the tree holes from which the population used in our experiment was sourced (black dots) as well as the locations of trapped *Ae. sierrensis* adults based on vector surveillance data from 2010 – 2020 obtained from CalSurv. Note this map does not depict the entire *Ae. sierrensis* range, which extends from approximately southern California to British Columbia (16). Colors denote the current (2020) daily mean temperatures in the spring—the larval activity period.

**Table S1.** Sample numbers used in the experiment and sequencing approaches. Note, analyses were conducted only on those samples with quality sequence data (n = 227). Survival is expressed as a proportion.

| Treatment | Round | Rep. | Total Larvae | Total Pupae | Total Adults | Used | Sex (F/M) | Larval Survival | Pupal Survival |
|---|---|---|---|---|---|---|---|---|---|
| Control | 1 | A | 37 | 34 | 34 | 16 | 13/3 | 0.92 | 1.00 |
| Control | 1 | B | 43 | 39 | 38 | 28 | 12/16 | 0.91 | 0.97 |
| Heat-selected | 1 | C | 78 | 34 | 21 | 8 | 5/3 | 0.44 | 0.62 |
| Heat-selected | 1 | D | 77 | 42 | 28 | 10 | 7/3 | 0.55 | 0.67 |
| Control | 2 | A | 23 | 9 | 8 | 1 | 0/1 | 0.39 | 0.89 |
| Control | 2 | B | 28 | 11 | 11 | 3 | 2/1 | 0.39 | 1.00 |
| Heat-selected | 2 | C | 44 | 9 | 5 | 0 | 0/0 | 0.20 | 0.56 |
| Heat-selected | 2 | D | 53 | 6 | 1 | 0 | 0/0 | 0.11 | 0.17 |
| Control | 3 | A | 349 | 193 | 179 | 34 | 16/18 | 0.55 | 0.93 |
| Control | 3 | B | 310 | 168 | 153 | 40 | 23/17 | 0.54 | 0.91 |
| Heat-selected | 3 | C | 866 | 158 | 117 | 38 | 16/22 | 0.18 | 0.74 |
| Heat-selected | 3 | D | 825 | 222 | 178 | 49 | 24/25 | 0.27 | 0.80 |

**Table S2.** Summary results from knockdown assay and wing length measurements by treatment and sex.

| | | Knockdown time (minutes) | | | Wing length (mm) | | |
|---|---|---|---|---|---|---|---|
| **Treatment** | **Sex** | **Mean** | **Min** | **Max** | **Mean** | **Min** | **Max** |
| control | F | 48.57 | 32.70 | 67.62 | 3.15 | 2.86 | 3.36 |
| | M | 50.63 | 33.38 | 66.45 | 2.59 | 2.36 | 3.10 |
| heat | F | 45.11 | 19.78 | 63.50 | 2.68 | 2.37 | 2.88 |
| | M | 47.70 | 21.82 | 64.82 | 2.37 | 220 | 2.55 |

**Table S3.** Results of linear regression of knockdown time against treatment (heat-selected as reference), sex (male as reference), and wing length. Note that experimental round and knockdown assay round were included as random effects in this model.

| | Coef. estimate | Std. error | t value | p |
|---|---|---|---|---|
| Intercept | 65.62 | 14.37 | 4.57 | <0.001 |
| Treatment | -4.23 | 1.95 | -2.16 | 0.032 |
| Sex | -1.37 | 2.39 | -0.57 | 0.558 |
| Wing length | -5.39 | 4.60 | -1.17 | 0.244 |

**Table S4.** Individual sample statistics on sequence alignment and diversity. Each sample here refers to an individual mosquito used in the experiment. Note that observed and expected heterozygosity are only provided for control individuals, as these more accurately reflect the starting population.

| Sample | Treatment | Total reads | % Mapped reads | Mean sequence depth | Observed Het. | Expected Het. |
|---|---|---|---|---|---|---|
| E-010 | control | 88972171 | 97.94% | 9.26 | 0.342 | 0.370 |
| E-011 | control | 91180800 | 98.47% | 9.57 | 0.400 | 0.370 |
| E-012 | control | 123080899 | 98.44% | 12.91 | 0.409 | 0.370 |
| E-013 | control | 106645608 | 98.66% | 11.14 | 0.417 | 0.370 |
| E-015 | control | 76974036 | 98.82% | 8.10 | 0.431 | 0.370 |
| E-018 | control | 74255290 | 98.83% | 7.80 | 0.401 | 0.370 |
| E-019 | control | 74088272 | 98.66% | 7.80 | 0.397 | 0.370 |
| E-020 | control | 61114834 | 98.67% | 6.42 | 0.339 | 0.370 |
| E-022 | control | 95228421 | 98.76% | 10.02 | 0.348 | 0.370 |
| E-023 | control | 102024988 | 98.73% | 10.63 | 0.400 | 0.370 |
| E-024 | control | 111264028 | 98.63% | 11.64 | 0.419 | 0.370 |
| E-025 | control | 111881403 | 98.49% | 11.62 | 0.479 | 0.370 |
| E-026 | control | 106231424 | 98.70% | 11.15 | 0.438 | 0.370 |
| E-027 | control | 119205429 | 98.62% | 12.45 | 0.421 | 0.370 |
| E-028 | control | 98331757 | 98.68% | 10.36 | 0.384 | 0.370 |
| E-04 | control | 105284477 | 98.30% | 11.03 | 0.375 | 0.370 |
| F-002 | control | 103226543 | 98.65% | 10.81 | 0.423 | 0.370 |
| F-003 | control | 90439343 | 98.73% | 9.48 | 0.392 | 0.370 |
| F-005 | control | 85619893 | 98.66% | 8.99 | 0.404 | 0.370 |
| F-009 | control | 81369966 | 98.65% | 8.54 | 0.383 | 0.370 |
| F-011 | control | 138721784 | 98.51% | 14.43 | 0.407 | 0.370 |
| F-012 | control | 100235903 | 98.73% | 10.48 | 0.412 | 0.370 |
| F-013 | control | 102963402 | 98.61% | 10.77 | 0.475 | 0.370 |
| F-015 | control | 84087649 | 98.79% | 8.86 | 0.347 | 0.370 |

| | | | | | | |
|---|---|---|---|---|---|---|
| F-016 | control | 105204907 | 98.80% | 11.04 | 0.431 | 0.370 |
| F-017 | control | 81546746 | 98.68% | 8.52 | 0.380 | 0.370 |
| F-018 | control | 86065383 | 98.70% | 9.06 | 0.374 | 0.370 |
| F-019 | control | 122704865 | 98.60% | 12.87 | 0.421 | 0.370 |
| F-020 | control | 89199855 | 98.69% | 9.33 | 0.427 | 0.370 |
| F-021 | control | 90796842 | 98.85% | 9.51 | 0.370 | 0.370 |
| F-022 | control | 77410540 | 98.87% | 8.11 | 0.404 | 0.370 |
| F-023 | control | 123741287 | 98.65% | 12.92 | 0.409 | 0.370 |
| F-024 | control | 87341268 | 98.73% | 9.12 | 0.376 | 0.370 |
| F-025 | control | 119833051 | 98.62% | 12.57 | 0.426 | 0.370 |
| F-026 | control | 105675748 | 98.55% | 11.06 | 0.411 | 0.370 |
| F-027 | control | 106465057 | 98.72% | 11.11 | 0.367 | 0.370 |
| F-028 | control | 66272989 | 98.68% | 6.95 | 0.370 | 0.370 |
| F-029 | control | 105051781 | 98.76% | 11.01 | 0.432 | 0.370 |
| F-031 | control | 92105766 | 98.80% | 9.65 | 0.424 | 0.370 |
| F-032 | control | 120995454 | 98.71% | 12.67 | 0.354 | 0.370 |
| F-035 | control | 76570617 | 98.81% | 8.00 | 0.419 | 0.370 |
| F-036 | control | 85334612 | 98.57% | 8.96 | 0.405 | 0.370 |
| F-037 | control | 78419091 | 97.92% | 7.89 | 0.362 | 0.370 |
| F-038 | control | 106989570 | 98.44% | 11.12 | 0.326 | 0.370 |
| I-001 | control | 124771585 | 98.38% | 13.03 | 0.454 | 0.370 |
| J-001 | control | 87829264 | 98.40% | 9.17 | 0.403 | 0.370 |
| J-003 | control | 107479880 | 98.43% | 11.25 | 0.458 | 0.370 |
| J-004 | | 90974115 | 98.49% | 9.51 | 0.461 | 0.371 |
| M-001 | control | 95881713 | 98.56% | 10.01 | 0.412 | 0.370 |
| M-002 | control | 98902279 | 98.37% | 10.33 | 0.377 | 0.370 |
| M-003 | control | 90568406 | 98.42% | 9.44 | 0.413 | 0.370 |
| M-004 | control | 106479562 | 98.63% | 11.16 | 0.412 | 0.370 |

| | | | | | | |
|---|---|---|---|---|---|---|
| M-005 | control | 104546938 | 98.59% | 10.93 | 0.370 | 0.370 |
| M-006 | control | 96308825 | 98.55% | 10.02 | 0.414 | 0.370 |
| M-007 | control | 84926924 | 98.22% | 8.84 | 0.389 | 0.370 |
| M-008 | control | 82360248 | 98.11% | 8.51 | 0.378 | 0.370 |
| M-009 | control | 94919883 | 98.42% | 9.94 | 0.374 | 0.370 |
| M-010 | control | 90511413 | 98.41% | 9.51 | 0.390 | 0.370 |
| M-011 | control | 124909674 | 98.51% | 13.10 | 0.419 | 0.370 |
| M-017 | control | 59744582 | 98.49% | 6.28 | 0.376 | 0.370 |
| M-018 | control | 71144193 | 98.44% | 7.46 | 0.323 | 0.370 |
| M-019 | control | 77910407 | 98.74% | 8.20 | 0.431 | 0.370 |
| M-020 | control | 86956394 | 98.46% | 9.07 | 0.417 | 0.370 |
| M-022 | control | 156321929 | 98.42% | 16.31 | 0.415 | 0.370 |
| M-023 | control | 90047176 | 98.13% | 9.37 | 0.370 | 0.370 |
| M-024 | control | 74674809 | 98.59% | 7.83 | 0.367 | 0.370 |
| M-025 | control | 82051619 | 98.35% | 8.62 | 0.390 | 0.370 |
| M-026 | control | 81781725 | 98.73% | 8.55 | 0.383 | 0.370 |
| M-027 | control | 97196200 | 98.62% | 10.22 | 0.386 | 0.370 |
| M-028 | control | 111288661 | 98.59% | 11.67 | 0.339 | 0.370 |
| M-029 | control | 102621502 | 98.36% | 10.71 | 0.312 | 0.370 |
| M-030 | control | 90741223 | 98.57% | 9.47 | 0.375 | 0.370 |
| M-032 | control | 72951088 | 98.61% | 7.66 | 0.312 | 0.370 |
| M-033 | control | 77134442 | 98.53% | 8.08 | 0.399 | 0.370 |
| M-034 | control | 95701717 | 98.56% | 10.05 | 0.437 | 0.370 |
| M-035 | control | 72142544 | 98.32% | 7.55 | 0.369 | 0.370 |
| M-036 | control | 71596388 | 98.59% | 7.48 | 0.394 | 0.370 |
| M-037 | control | 60118290 | 98.57% | 6.29 | 0.408 | 0.370 |
| M-038 | control | 66764093 | 98.53% | 6.98 | 0.375 | 0.370 |
| M-039 | control | 102423830 | 98.16% | 10.66 | 0.414 | 0.370 |

| | | | | | | |
|---|---|---|---|---|---|---|
| M-040 | control | 96750799 | 98.33% | 9.97 | 0.356 | 0.370 |
| M-041 | control | 135236014 | 98.43% | 14.06 | 0.390 | 0.370 |
| N-001 | control | 101746192 | 98.41% | 10.55 | 0.429 | 0.370 |
| N-002 | control | 92154762 | 98.64% | 9.63 | 0.419 | 0.370 |
| N-003 | control | 111229942 | 98.56% | 11.59 | 0.426 | 0.370 |
| N-004 | control | 145629476 | 98.55% | 15.21 | 0.436 | 0.370 |
| N-005 | control | 147250436 | 98.68% | 15.35 | 0.428 | 0.370 |
| N-006 | control | 90022696 | 98.26% | 9.36 | 0.400 | 0.370 |
| N-007 | control | 76935794 | 98.40% | 8.01 | 0.377 | 0.370 |
| N-008 | control | 97212880 | 98.58% | 10.14 | 0.382 | 0.370 |
| N-009 | control | 114113508 | 98.58% | 11.90 | 0.430 | 0.370 |
| N-010 | control | 83756800 | 98.38% | 8.66 | 0.417 | 0.370 |
| N-011 | control | 95867382 | 98.12% | 9.92 | 0.392 | 0.370 |
| N-012 | control | 111884451 | 97.97% | 11.57 | 0.424 | 0.370 |
| N-013 | control | 89151847 | 98.29% | 9.21 | 0.394 | 0.370 |
| N-018 | control | 83972464 | 98.04% | 8.67 | 0.416 | 0.370 |
| N-019 | control | 93335110 | 98.16% | 9.70 | 0.370 | 0.370 |
| N-020 | control | 90184215 | 97.93% | 9.29 | 0.421 | 0.370 |
| N-021 | control | 74920348 | 97.98% | 7.77 | 0.352 | 0.370 |
| N-022 | control | 91630730 | 97.55% | 9.36 | 0.349 | 0.370 |
| N-023 | control | 81569877 | 97.98% | 8.44 | 0.391 | 0.370 |
| N-024 | control | 68791313 | 98.12% | 7.14 | 0.377 | 0.370 |
| N-025 | control | 74686600 | 98.04% | 7.74 | 0.359 | 0.370 |
| N-026 | control | 120401744 | 97.93% | 12.45 | 0.333 | 0.370 |
| N-027 | control | 102529265 | 97.78% | 10.58 | 0.419 | 0.370 |
| N-028 | control | 113604225 | 98.04% | 11.75 | 0.437 | 0.370 |
| N-029 | control | 87042055 | 98.07% | 9.04 | 0.401 | 0.370 |
| N-030 | control | 98718443 | 98.11% | 10.21 | 0.412 | 0.370 |

| | | | | | | |
|---|---|---|---|---|---|---|
| N-032 | control | 73527401 | 98.24% | 7.65 | 0.401 | 0.370 |
| N-033 | control | 86051142 | 97.97% | 8.93 | 0.331 | 0.370 |
| N-034 | control | 90672674 | 97.85% | 9.34 | 0.411 | 0.370 |
| N-035 | control | 93743134 | 98.20% | 9.71 | 0.376 | 0.370 |
| N-036 | control | 90115053 | 98.17% | 9.33 | 0.309 | 0.370 |
| N-037 | control | 101246270 | 98.18% | 10.49 | 0.415 | 0.370 |
| N-038 | control | 84131705 | 98.13% | 8.71 | 0.371 | 0.370 |
| N-039 | control | 104797916 | 98.11% | 10.91 | 0.362 | 0.370 |
| N-040 | control | 118441947 | 98.22% | 12.28 | 0.427 | 0.370 |
| N-041 | control | 88055052 | 97.84% | 9.06 | 0.323 | 0.370 |
| N-042 | control | 112354015 | 98.28% | 11.60 | 0.381 | 0.370 |
| N-043 | control | 77002866 | 98.04% | 7.98 | 0.355 | 0.370 |
| N-044 | control | 98944175 | 98.13% | 10.25 | 0.347 | 0.370 |
| N-045 | control | 133004594 | 98.04% | 13.79 | 0.414 | 0.370 |
| O-003 | heat-selected | 87489748 | 97.84% | 9.06 | | |
| G-003 | heat-selected | 124882679 | 98.43% | 13.06 | | |
| G-004 | heat-selected | 70691083 | 98.57% | 7.41 | | |
| G-007 | heat-selected | 151424539 | 98.64% | 15.90 | | |
| G-016 | heat-selected | 58541530 | 99.00% | 5.99 | | |
| G-017 | heat-selected | 112571090 | 98.98% | 11.66 | | |
| G-018 | heat-selected | 88295251 | 98.79% | 9.14 | | |
| G-019 | heat-selected | 126428655 | 98.42% | 13.28 | | |
| G-020 | heat-selected | 105416004 | 98.45% | 11.08 | | |
| H-005 | heat-selected | 113264831 | 98.45% | 11.92 | | |
| H-006 | heat-selected | 95601389 | 98.13% | 9.98 | | |
| H-007 | heat-selected | 128167590 | 98.23% | 13.43 | | |
| H-008 | heat-selected | 105755875 | 98.28% | 11.07 | | |
| H-010 | heat-selected | 99082242 | 98.45% | 10.37 | | |

| | | | | |
|---|---|---|---|---|
| H-020 | heat-selected | 87087854 | 98.22% | 9.11 |
| H-021 | heat-selected | 99204662 | 98.03% | 10.33 |
| H-022 | heat-selected | 113404177 | 98.58% | 11.81 |
| H-023 | heat-selected | 101521940 | 98.66% | 10.66 |
| H-025 | heat-selected | 101240619 | 98.76% | 10.62 |
| O-001 | heat-selected | 64885478 | 98.23% | 6.73 |
| O-002 | heat-selected | 108690870 | 98.13% | 11.25 |
| O-004 | heat-selected | 90813271 | 98.14% | 9.40 |
| O-005 | heat-selected | 95898538 | 97.90% | 9.88 |
| O-006 | heat-selected | 100423660 | 98.02% | 10.34 |
| O-007 | heat-selected | 106289300 | 98.08% | 10.98 |
| O-008 | heat-selected | 121659828 | 97.83% | 12.51 |
| O-009 | heat-selected | 102138421 | 98.04% | 10.58 |
| O-010 | heat-selected | 98215117 | 98.11% | 10.11 |
| O-011 | heat-selected | 72220572 | 98.16% | 7.47 |
| O-012 | heat-selected | 100107368 | 98.26% | 10.29 |
| O-013 | heat-selected | 87816035 | 98.02% | 9.06 |
| O-015 | heat-selected | 93749441 | 98.47% | 9.64 |
| O-016 | heat-selected | 90816467 | 98.33% | 9.41 |
| O-017 | heat-selected | 83627170 | 98.16% | 8.65 |
| O-018 | heat-selected | 79626884 | 97.54% | 8.17 |
| O-019 | heat-selected | 115157546 | 98.17% | 11.92 |
| O-020 | heat-selected | 98482767 | 98.50% | 10.23 |
| O-022 | heat-selected | 82643180 | 98.18% | 8.57 |
| O-023 | heat-selected | 103798890 | 98.43% | 10.82 |
| O-024 | heat-selected | 92680918 | 98.02% | 9.63 |
| O-025 | heat-selected | 94620439 | 98.40% | 9.87 |
| O-026 | heat-selected | 115620795 | 98.23% | 12.02 |

| | | | | |
|---|---|---|---|---|
| O-027 | heat-selected | 133123223 | 98.19% | 13.84 |
| O-028 | heat-selected | 80327388 | 98.03% | 8.36 |
| O-030 | heat-selected | 93801698 | 97.94% | 9.72 |
| O-031 | heat-selected | 84362504 | 98.16% | 8.76 |
| O-032 | heat-selected | 103207027 | 98.49% | 10.77 |
| O-033 | heat-selected | 80051186 | 98.14% | 8.28 |
| O-034 | heat-selected | 116313575 | 98.42% | 12.13 |
| O-035 | heat-selected | 116949525 | 98.64% | 12.16 |
| O-036 | heat-selected | 104366833 | 98.37% | 10.79 |
| O-037 | heat-selected | 84254135 | 98.30% | 8.74 |
| O-038 | heat-selected | 68948164 | 98.38% | 7.21 |
| O-039 | heat-selected | 82959014 | 98.71% | 8.65 |
| O-040 | heat-selected | 58640191 | 98.20% | 6.13 |
| O-041 | heat-selected | 99058381 | 98.39% | 10.34 |
| P-001 | heat-selected | 80627305 | 98.44% | 8.40 |
| P-002 | heat-selected | 80048623 | 98.44% | 8.22 |
| P-003 | heat-selected | 84377421 | 98.17% | 8.74 |
| P-004 | heat-selected | 88228667 | 98.40% | 9.20 |
| P-005 | heat-selected | 76453813 | 98.45% | 7.98 |
| P-006 | heat-selected | 83886813 | 98.28% | 8.72 |
| P-007 | heat-selected | 123843743 | 98.47% | 12.80 |
| P-008 | heat-selected | 85407210 | 98.46% | 8.84 |
| P-009 | heat-selected | 83196975 | 98.68% | 8.60 |
| P-010 | heat-selected | 113997019 | 98.26% | 11.83 |
| P-011 | heat-selected | 77707534 | 98.29% | 8.10 |
| P-012 | heat-selected | 117111546 | 98.40% | 12.17 |
| P-013 | heat-selected | 125661302 | 98.49% | 13.06 |
| P-014 | heat-selected | 92988212 | 98.68% | 9.65 |

| | | | | |
|---|---|---|---|---|
| P-015 | heat-selected | 61639393 | 98.37% | 6.45 |
| P-016 | heat-selected | 109799039 | 98.31% | 11.43 |
| P-017 | heat-selected | 170831688 | 98.52% | 17.83 |
| P-018 | heat-selected | 93632415 | 98.20% | 9.74 |
| P-019 | heat-selected | 133279867 | 98.36% | 13.89 |
| P-020 | heat-selected | 140287750 | 98.57% | 14.67 |
| P-021 | heat-selected | 101250906 | 98.28% | 10.55 |
| P-022 | heat-selected | 107617026 | 98.24% | 11.19 |
| P-023 | heat-selected | 111406762 | 98.26% | 11.61 |
| P-024 | heat-selected | 70473618 | 97.93% | 7.26 |
| P-025 | heat-selected | 79050704 | 98.23% | 8.18 |
| P-026 | heat-selected | 86637790 | 98.26% | 8.98 |
| P-027 | heat-selected | 68037819 | 98.22% | 7.10 |
| P-028 | heat-selected | 99498922 | 98.18% | 10.28 |
| P-029 | heat-selected | 100802264 | 98.11% | 10.47 |
| P-030 | heat-selected | 114314058 | 98.09% | 11.88 |
| P-031 | heat-selected | 93148313 | 97.80% | 9.66 |
| P-032 | heat-selected | 107567583 | 98.14% | 11.21 |
| P-033 | heat-selected | 87975325 | 98.46% | 9.20 |
| P-034 | heat-selected | 87033212 | 98.17% | 9.03 |
| P-035 | heat-selected | 87475653 | 97.96% | 9.08 |
| P-036 | heat-selected | 88895339 | 98.37% | 9.26 |
| P-037 | heat-selected | 67692522 | 98.06% | 7.03 |
| P-038 | heat-selected | 77601487 | 98.18% | 8.06 |
| P-039 | heat-selected | 96829478 | 98.09% | 10.09 |
| P-040 | heat-selected | 90311837 | 97.92% | 9.41 |
| P-041 | heat-selected | 72032041 | 98.30% | 7.52 |
| P-042 | heat-selected | 89089536 | 98.41% | 9.28 |

| | | | | |
|---|---|---|---|---|
| P-044 | heat-selected | 85540404 | 98.48% | 8.96 |
| P-045 | heat-selected | 96453611 | 98.11% | 10.05 |
| P-046 | heat-selected | 82208548 | 98.08% | 8.54 |
| P-047 | heat-selected | 51703633 | 97.98% | 5.38 |
| P-048 | heat-selected | 63535808 | 98.17% | 6.64 |
| P-049 | heat-selected | 93541975 | 98.19% | 9.74 |
| P-050 | heat-selected | 86659420 | 97.98% | 8.99 |

**Table S5.** Individual sample metadata and experimental results. Herein, 'Sample' is a unique label assigned to an individual mosquito used in the experiment, 'Time' denotes the number of hours since adult eclosion, and 'Round' refers to the experimental round (1-3).

| Sample | Treatment | Replicate | Sex | Time | Round | Wing length (mm) | Knockdown time (mins) |
|---|---|---|---|---|---|---|---|
| E-010 | control | E | F | 48h | 1 | 3.15 | 43.08 |
| E-011 | control | E | F | 48h | 1 | 3.16 | 59.40 |
| E-012 | control | E | F | 48h | 1 | 3.09 | 59.95 |
| E-013 | control | E | M | 48h | 1 | 2.57 | 54.97 |
| E-015 | control | E | F | 48h | 1 | 3.12 | 58.33 |
| E-018 | control | E | F | 48h | 1 | 3.23 | 49.92 |
| E-019 | control | E | F | 48h | 1 | 3.04 | 51.57 |
| E-020 | control | E | F | 48h | 1 | 3.32 | 47.52 |
| E-022 | control | E | M | 48h | 1 | 3.12 | 44.67 |
| E-023 | control | E | F | 48h | 1 | 3.13 | 57.00 |
| E-024 | control | E | F | 48h | 1 | 3.06 | 46.88 |
| E-025 | control | E | F | 48h | 1 | NA | 58.25 |
| E-026 | control | E | F | 48h | 1 | 3.13 | 45.33 |
| E-027 | control | E | F | 48h | 1 | NA | 40.92 |
| E-028 | control | E | F | 48h | 1 | 3.22 | 41.10 |
| E-04 | control | E | M | 48h | 1 | 2.63 | 57.63 |
| F-002 | control | F | M | 48h | 1 | 2.65 | 54.73 |
| F-003 | control | F | M | 48h | 1 | NA | 66.10 |
| F-005 | control | F | M | 48h | 1 | 2.64 | 66.45 |
| F-009 | control | F | M | 48h | 1 | 2.66 | 55.30 |
| F-011 | control | F | M | 48h | 1 | NA | 51.47 |
| F-012 | control | F | M | 48h | 1 | 2.71 | 59.05 |
| F-013 | control | F | M | 48h | 1 | NA | 47.37 |
| F-015 | control | F | M | 48h | 1 | 2.62 | 57.50 |
| F-016 | control | F | F | 48h | 1 | 3.23 | 47.88 |
| F-017 | control | F | M | 48h | 1 | 2.53 | 51.73 |
| F-018 | control | F | F | 48h | 1 | 3.18 | 54.38 |
| F-019 | control | F | F | 48h | 1 | 3.04 | 67.47 |
| F-020 | control | F | M | 48h | 1 | 2.61 | 47.62 |
| F-021 | control | F | M | 48h | 1 | 2.66 | 54.42 |
| F-022 | control | F | F | 48h | 1 | NA | 52.80 |
| F-023 | control | F | M | 48h | 1 | 2.70 | 48.22 |
| F-024 | control | F | M | 48h | 1 | NA | 63.33 |
| F-025 | control | F | F | 48h | 1 | NA | 49.03 |
| F-026 | control | F | M | 48h | 1 | NA | 48.58 |

| | | | | | | | |
|---|---|---|---|---|---|---|---|
| F-027 | control | F | M | 48h | 1 | NA | 47.62 |
| F-028 | control | F | F | 48h | 1 | 3.24 | 54.97 |
| F-029 | control | F | F | 48h | 1 | 3.12 | 41.92 |
| F-031 | control | F | F | 48h | 1 | 3.18 | 43.47 |
| F-032 | control | F | F | 48h | 1 | 3.13 | 57.38 |
| F-035 | control | F | F | 48h | 1 | 3.20 | 50.65 |
| F-036 | control | F | F | 48h | 1 | 3.22 | 39.67 |
| F-037 | control | F | M | 48h | 1 | 2.69 | 44.23 |
| F-038 | control | F | F | 48h | 1 | 3.22 | 41.73 |
| I-001 | control | I | M | 48h | 2 | 2.54 | 43.97 |
| J-001 | control | J | M | 48h | 2 | NA | 58.33 |
| J-003 | control | J | F | 48h | 2 | 3.07 | 44.03 |
| M-001 | control | M | M | 48h | 3 | 2.56 | 61.15 |
| M-002 | control | M | M | 48h | 3 | NA | 41.33 |
| M-003 | control | M | M | 48h | 3 | NA | 46.67 |
| M-004 | control | M | F | 48h | 3 | 2.97 | 67.62 |
| M-005 | control | M | M | 48h | 3 | 2.66 | 59.50 |
| M-006 | control | M | M | 48h | 3 | 2.66 | 59.83 |
| M-007 | control | M | F | 72h | 3 | 3.05 | 50.82 |
| M-008 | control | M | M | 72h | 3 | 2.50 | 58.50 |
| M-009 | control | M | M | 72h | 3 | 2.56 | 51.42 |
| M-010 | control | M | M | 72h | 3 | 2.52 | 41.62 |
| M-011 | control | M | M | 72h | 3 | 2.57 | 44.95 |
| M-017 | control | M | F | 72h | 3 | 3.08 | 48.73 |
| M-018 | control | M | M | 72h | 3 | 2.40 | 51.72 |
| M-019 | control | M | F | 72h | 3 | 3.06 | 48.35 |
| M-020 | control | M | M | 72h | 3 | 2.61 | 39.28 |
| M-022 | control | M | M | 72h | 3 | 2.59 | 56.88 |
| M-023 | control | M | F | 72h | 3 | 3.27 | 50.50 |
| M-024 | control | M | M | 72h | 3 | 2.60 | 40.20 |
| M-025 | control | M | F | 72h | 3 | 3.04 | 46.93 |
| M-026 | control | M | M | 72h | 3 | NA | 55.93 |
| M-027 | control | M | F | 72h | 3 | 3.13 | 50.50 |
| M-028 | control | M | M | 72h | 3 | 3.09 | 41.77 |
| M-029 | control | M | F | 72h | 3 | 3.17 | 55.45 |
| M-030 | control | M | M | 72h | 3 | 2.81 | 33.38 |
| M-032 | control | M | M | 72h | 3 | NA | 41.42 |
| M-033 | control | M | F | 72h | 3 | 3.19 | 42.13 |
| M-034 | control | M | F | 72h | 3 | 3.22 | 56.80 |

| | | | | | | | |
|---|---|---|---|---|---|---|---|
| M-035 | control | M | F | 72h | 3 | 3.20 | 51.58 |
| M-036 | control | M | F | 72h | 3 | NA | 50.27 |
| M-037 | control | M | F | 72h | 3 | 3.08 | 43.23 |
| M-038 | control | M | F | 72h | 3 | 2.86 | 33.53 |
| M-039 | control | M | F | 72h | 3 | 3.11 | 44.42 |
| M-040 | control | M | M | 72h | 3 | 2.54 | 48.77 |
| M-041 | control | M | F | 72h | 3 | 3.21 | 55.72 |
| N-001 | control | N | M | 48h | 3 | 2.47 | 43.80 |
| N-002 | control | N | M | 48h | 3 | NA | 45.62 |
| N-003 | control | N | M | 48h | 3 | 2.67 | 50.83 |
| N-004 | control | N | F | 48h | 3 | NA | 45.05 |
| N-005 | control | N | M | 48h | 3 | NA | 61.53 |
| N-006 | control | N | F | 48h | 3 | NA | 50.37 |
| N-007 | control | N | M | 48h | 3 | NA | 45.85 |
| N-008 | control | N | M | 72h | 3 | 2.60 | 48.13 |
| N-009 | control | N | M | 72h | 3 | 2.44 | 41.37 |
| N-010 | control | N | M | 72h | 3 | 2.62 | 52.48 |
| N-011 | control | N | M | 72h | 3 | 2.57 | 50.20 |
| N-012 | control | N | M | 72h | 3 | 2.41 | 51.37 |
| N-013 | control | N | M | 72h | 3 | 2.36 | 47.47 |
| N-018 | control | N | M | 72h | 3 | 2.52 | 39.93 |
| N-019 | control | N | F | 72h | 3 | 3.14 | 52.67 |
| N-020 | control | N | M | 72h | 3 | 2.65 | 49.85 |
| N-021 | control | N | F | 72h | 3 | 3.27 | 46.10 |
| N-022 | control | N | M | 72h | 3 | 2.51 | 43.98 |
| N-023 | control | N | F | 72h | 3 | NA | 32.70 |
| N-024 | control | N | M | 72h | 3 | NA | 52.83 |
| N-025 | control | N | F | 72h | 3 | NA | 39.00 |
| N-026 | control | N | M | 72h | 3 | 2.42 | 37.53 |
| N-027 | control | N | F | 72h | 3 | 3.26 | 54.27 |
| N-028 | control | N | M | 72h | 3 | 2.55 | 55.02 |
| N-029 | control | N | F | 72h | 3 | NA | 53.77 |
| N-030 | control | N | M | 72h | 3 | 2.53 | 59.48 |
| N-032 | control | N | M | 72h | 3 | 2.50 | 52.03 |
| N-033 | control | N | F | 72h | 3 | 3.24 | 44.33 |
| N-034 | control | N | M | 72h | 3 | 2.60 | 44.63 |
| N-035 | control | N | F | 72h | 3 | 3.19 | 36.88 |
| N-036 | control | N | M | 72h | 3 | 2.43 | 63.55 |
| N-037 | control | N | F | 72h | 3 | 3.24 | 39.53 |

| | | | | | | | |
|---|---|---|---|---|---|---|---|
| N-038 | control | N | M | 72h | 3 | 2.71 | 47.93 |
| N-039 | control | N | F | 72h | 3 | 3.36 | 39.30 |
| N-040 | control | N | F | 72h | 3 | NA | 42.80 |
| N-041 | control | N | F | 72h | 3 | 3.03 | 47.50 |
| N-042 | control | N | M | 72h | 3 | 2.30 | 57.30 |
| N-043 | control | N | F | 72h | 3 | 2.58 | 45.30 |
| N-044 | control | N | F | 72h | 3 | 2.91 | 45.88 |
| N-045 | control | N | F | 72h | 3 | 3.35 | 44.53 |
| O-003 | control | O | M | 48h | 3 | NA | 59.10 |
| G-003 | heat | G | F | 48h | 1 | NA | 41.85 |
| G-004 | heat | G | F | 48h | 1 | 2.37 | 52.33 |
| G-007 | heat | G | M | 48h | 1 | 2.52 | 58.82 |
| G-016 | heat | G | F | 48h | 1 | 2.70 | 47.30 |
| G-017 | heat | G | F | 48h | 1 | NA | 56.25 |
| G-018 | heat | G | M | 48h | 1 | 2.37 | 53.20 |
| G-019 | heat | G | F | 48h | 1 | 2.61 | 53.83 |
| G-020 | heat | G | M | 48h | 1 | 2.44 | 43.30 |
| H-005 | heat | H | F | 48h | 1 | 2.70 | 52.57 |
| H-006 | heat | H | F | 48h | 1 | 2.77 | 60.77 |
| H-007 | heat | H | F | 48h | 1 | 2.70 | 58.45 |
| H-008 | heat | H | M | 48h | 1 | 2.49 | 42.83 |
| H-010 | heat | H | F | 48h | 1 | 2.80 | 51.25 |
| H-020 | heat | H | M | 48h | 1 | 2.22 | 42.02 |
| H-021 | heat | H | F | 48h | 1 | NA | 54.83 |
| H-022 | heat | H | M | 48h | 1 | 2.25 | 42.63 |
| H-023 | heat | H | F | 48h | 1 | NA | 44.57 |
| H-025 | heat | H | F | 48h | 1 | 2.37 | 44.75 |
| J-004 | heat | J | F | 48h | 2 | NA | 34.12 |
| O-001 | heat | O | M | 48h | 3 | NA | 54.43 |
| O-002 | heat | O | M | 48h | 3 | 2.33 | 59.42 |
| O-004 | heat | O | M | 48h | 3 | 2.46 | 54.60 |
| O-005 | heat | O | M | 48h | 3 | 2.28 | 49.53 |
| O-006 | heat | O | M | 72h | 3 | NA | 61.77 |
| O-007 | heat | O | M | 72h | 3 | 2.33 | 46.22 |
| O-008 | heat | O | M | 72h | 3 | 2.31 | 57.03 |
| O-009 | heat | O | M | 72h | 3 | NA | 40.42 |
| O-010 | heat | O | M | 72h | 3 | NA | 40.98 |
| O-011 | heat | O | M | 72h | 3 | NA | 58.58 |
| O-012 | heat | O | M | 72h | 3 | 2.36 | 43.42 |

| | | | | | | | |
|---|---|---|---|---|---|---|---|
| O-013 | heat | O | F | 72h | 3 | 2.84 | 41.45 |
| O-015 | heat | O | F | 72h | 3 | NA | 50.50 |
| O-016 | heat | O | M | 72h | 3 | NA | 47.23 |
| O-017 | heat | O | F | 72h | 3 | 2.87 | 43.50 |
| O-018 | heat | O | M | 72h | 3 | 2.35 | 41.50 |
| O-019 | heat | O | M | 72h | 3 | NA | 42.02 |
| O-020 | heat | O | F | 72h | 3 | 2.64 | 42.90 |
| O-022 | heat | O | F | 72h | 3 | 2.88 | 51.25 |
| O-023 | heat | O | M | 72h | 3 | 2.30 | 50.87 |
| O-024 | heat | O | F | 72h | 3 | 2.78 | 42.78 |
| O-025 | heat | O | M | 72h | 3 | 2.20 | 59.38 |
| O-026 | heat | O | F | 72h | 3 | 2.45 | 49.63 |
| O-027 | heat | O | M | 72h | 3 | 2.33 | 40.62 |
| O-028 | heat | O | F | 72h | 3 | 2.56 | 44.00 |
| O-030 | heat | O | F | 72h | 3 | 2.62 | 53.02 |
| O-031 | heat | O | M | 72h | 3 | NA | 38.00 |
| O-032 | heat | O | F | 72h | 3 | NA | 47.77 |
| O-033 | heat | O | M | 72h | 3 | 2.25 | 38.40 |
| O-034 | heat | O | F | 72h | 3 | 2.72 | 53.47 |
| O-035 | heat | O | M | 72h | 3 | 2.34 | 47.13 |
| O-036 | heat | O | F | 72h | 3 | 2.49 | 48.65 |
| O-037 | heat | O | M | 72h | 3 | NA | 49.75 |
| O-038 | heat | O | F | 72h | 3 | 2.69 | 38.43 |
| O-039 | heat | O | F | 72h | 3 | 2.59 | 36.95 |
| O-040 | heat | O | F | 72h | 3 | NA | 49.80 |
| O-041 | heat | O | F | 72h | 3 | 2.31 | 63.50 |
| P-001 | heat | I | M | 48h | 3 | 2.38 | 43.93 |
| P-002 | heat | P | M | 48h | 3 | 2.43 | 49.48 |
| P-003 | heat | P | M | 72h | 3 | NA | 44.97 |
| P-004 | heat | P | M | 48h | 3 | NA | 63.58 |
| P-005 | heat | P | M | 72h | 3 | 2.37 | 52.83 |
| P-006 | heat | I | M | 72h | 3 | 2.55 | 47.50 |
| P-007 | heat | P | M | 72h | 3 | 2.41 | 51.98 |
| P-008 | heat | P | M | 72h | 3 | NA | 54.75 |
| P-009 | heat | P | F | 48h | 3 | 2.64 | 44.02 |
| P-010 | heat | P | M | 48h | 3 | 2.35 | 41.60 |
| P-011 | heat | P | F | 48h | 3 | 2.59 | 43.40 |
| P-012 | heat | P | F | 48h | 3 | 2.72 | 51.50 |
| P-013 | heat | P | F | 48h | 3 | 2.81 | 41.38 |

| | | | | | | | |
|---|---|---|---|---|---|---|---|
| P-014 | heat | P | F | 72h | 3 | 2.77 | 42.47 |
| P-015 | heat | P | M | 72h | 3 | 2.45 | 62.37 |
| P-016 | heat | P | M | 72h | 3 | 2.43 | 45.70 |
| P-017 | heat | P | M | 72h | 3 | NA | 41.52 |
| P-018 | heat | P | M | 72h | 3 | NA | 52.07 |
| P-019 | heat | P | M | 72h | 3 | NA | 64.82 |
| P-020 | heat | P | M | 72h | 3 | 2.49 | 50.02 |
| P-021 | heat | P | F | 72h | 3 | 2.78 | 41.98 |
| P-022 | heat | P | M | 72h | 3 | 2.36 | 45.03 |
| P-023 | heat | P | F | 72h | 3 | 2.70 | 36.28 |
| P-024 | heat | P | M | 72h | 3 | 2.48 | 41.07 |
| P-025 | heat | P | F | 72h | 3 | 2.73 | 44.28 |
| P-026 | heat | P | M | 72h | 3 | 2.46 | 35.23 |
| P-027 | heat | P | F | 72h | 3 | 2.68 | 49.12 |
| P-028 | heat | P | M | 72h | 3 | 2.29 | 30.53 |
| P-029 | heat | P | F | 72h | 3 | 2.73 | 38.35 |
| P-030 | heat | P | M | 72h | 3 | 2.25 | 25.40 |
| P-031 | heat | P | F | 72h | 3 | NA | 39.95 |
| P-032 | heat | P | M | 72h | 3 | 2.47 | 45.37 |
| P-033 | heat | P | F | 72h | 3 | 2.75 | 49.40 |
| P-034 | heat | P | M | 72h | 3 | NA | 47.33 |
| P-035 | heat | P | F | 72h | 3 | 2.72 | 32.02 |
| P-036 | heat | P | M | 72h | 3 | 2.32 | 58.00 |
| P-037 | heat | P | F | 72h | 3 | NA | 36.38 |
| P-038 | heat | P | M | 72h | 3 | 2.45 | 47.78 |
| P-039 | heat | P | F | 72h | 3 | NA | 47.05 |
| P-040 | heat | P | F | 72h | 3 | 2.53 | 59.82 |
| P-041 | heat | P | F | 72h | 3 | 2.47 | 34.90 |
| P-042 | heat | P | F | 72h | 3 | 2.86 | 51.58 |
| P-044 | heat | P | F | 72h | 3 | 2.69 | 40.95 |
| P-045 | heat | P | F | 72h | 3 | 2.47 | 39.28 |
| P-046 | heat | P | M | 72h | 3 | 2.34 | 21.82 |
| P-047 | heat | P | F | 72h | 3 | 2.47 | 19.78 |
| P-048 | heat | P | F | 72h | 3 | 2.52 | 33.05 |
| P-049 | heat | P | F | 72h | 3 | NA | 42.28 |
| P-050 | heat | P | F | 72h | 3 | NA | 20.87 |

**Table S6.** Characteristics of inversions that were significantly differentiated ($p < 0.05$ after Bonferroni correction for multiple testing) between control and heat-selected individuals or between individuals from the top 25% versus bottom 25% of knockdown times. The 'ROI' column denotes whether the inversion fell within the region of interest—the chromosomal regions in which there was an elevated number of SNPs associated with prolonged, larval heat tolerance (Figure 3A, Supplemental Methods).

| Inversions ID | Chr. | Start | End | Length | Frequency in control | Frequency in heat | ROI (T/F) |
|---|---|---|---|---|---|---|---|
| 0_1_INV00047889 | 1 | 243420308 | 312327110 | 68906801 | 13.3 | 4.9 | F |
| 20_1_INV00048775 | 1 | 274190042 | 276368387 | 2178344 | 2.0 | 6.9 | F |
| 13_1_INV00114975 | 2 | 117780421 | 193554100 | 75773678 | 5.4 | 0.5 | F |
| 8_1_INV00157996 | 2 | 359879989 | 428232800 | 68352810 | 11.8 | 3.4 | F |
| 0_1_INV00260322 | 3 | 197759320 | 373487690 | 175728369 | 14.8 | 5.4 | F |
| 5_1_INV00199156 | 3 | 266407859 | 370648399 | 104240539 | 13.3 | 3.9 | F |
| 1_1_INV00145236 | 2 | 244417907 | 257566837 | 13148929 | 8.9 | 1.5 | T |
| 1_1_INV00149242 | 2 | 258618054 | 275672360 | 17054305 | 5.9 | 1.0 | T |
| 0_1_INV00230929 | 3 | 112872175 | 117382970 | 4510794 | 3.0 | 7.9 | T |
| **Inversions ID** | **Chr.** | **Start** | **End** | **Length** | **Frequency in bottom 25%** | **Frequency in top 25%** | **ROI (T/F)** |
| 0_1_INV00044436 | 1 | 220680604 | 294626593 | 73945988 | 3.0 | 8.4 | F |
| 4_1_INV00103587 | 2 | 246541219 | 358808212 | 112266992 | 1.5 | 5.4 | T |
| 2_1_INV00143633 | 2 | 250046161 | 444498783 | 194452621 | 11.3 | 15.8 | T |
| 4_1_INV00143280 | 3 | 15864580 | 60675630 | 44811049 | 1.5 | 5.4 | T |
| 9_1_INV00178653 | 3 | 21007270 | 84943393 | 63936122 | 1.5 | 5.4 | T |

**Table S7.** Annotations for candidate genes associated with heat tolerance. The candidate genes listed below are those identified by two approaches. Here, '1' denotes the $F_{st}$ on treatment approach, '2' denotes the case-control GWA, and '3' denotes the GWA on knockdown time. Gene IDs are those assigned by *AUGUSTUS* or *GENEMARK-ES/ET*, while the NCBI accessions and Gene Names are based on the top BLASTN result. 'Chrom.' denotes the chromosomal position. 'NA's in Gene Names indicate genes lacking annotation/characterization in related species.

| Gene ID | Chrom. | Approach | NCBI accession | Gene Name | Relevant Prior Findings & Associations |
|---|---|---|---|---|---|
| g109 | 1 | 1 & 2 | XR_009997746.1 | NA | |
| g13206 | 2 | 1 & 2 | XR_009997695.1 | profilin | potential protective effect under heat stress, function as molecular chaperones (17, 18) |
| g13275 | 2 | 1 & 2 | NA | NA | |
| g13368 | 2 | 1 & 2 | XR_002502849.1 | NA | |
| g13376 | 2 | 1 & 2 | XM_029868494.2 | histone H3 | heat shock memory, enhanced survival under subsequent heat exposure (19, 20) |
| g13417 | 2 | 1 & 2 | XM_029880119.2 | NA | |
| gene_id file_2_17112_g | 2 | 1 & 2 | XM_062854850.1 | probable cytochrome P450 6a14 | environmental stress response (21–24) |
| g13524 | 2 | 1 & 2 | XM_029857670.2 | hemicentin-1 | environmental stress response, response to DENV infection (25, 26) |
| g13534 | 2 | 1 & 2 | XM_062860084.1 | uncharacterized | |
| g13617 | 2 | 1 & 2 | XM_062849755.1 | neural-cadherin | cell survival (27) |
| g13737 | 2 | 1 & 2 | XM_021844107.1 | sex peptide receptor | post-mating responses in *Ae. aegypti* (28) |
| g20424 | 3 | 1 & 2 | NA | NA | |
| g20752 | 3 | 1 & 2 | NA | NA | |
| g13490 | 2 | 1 & 3 | XM_019694398.3 | 2-phosphoxylose phosphatase 1 | |
| g22072 | 3 | 1 & 3 | XM_062845887.1 | NA | |
| g22315 | 3 | 1 & 3 | XM_021850513.1 | DNA repair endonuclease XPF | heat tolerance, desiccation resistance, interaction with heat shock proteins (29–31) |
| g13375 | 2 | 2 & 3 | XM_021844736.1 | nuclear pore complex protein Nup98-Nup96 | |
| g14290 | 2 | 2 & 3 | XM_062846923.1 | dopamine receptor 1 | olfactory learning in *Ae. aegypti* (32) |
| g22256 | 3 | 2 & 3 | FJ387159.1 | sensory neuron membrane protein 2 | odor perception in *Ae. albopictus* (33) |

**Table S8**. Parameter values used in the evolutionary rescue model to estimate adaptive potential. See Table S9 for values of rates of warming used in comparison to $\eta_c$, the maximum rate of environmental change under which populations could persist.

| Parameter | Estimate(s) | Source / Method |
|---|---|---|
| $r_{max}$ (max. population growth rate) | 0.187, 0.335, 0.379 | (34) Values are for *Anopheles, Aedes*, and *Culex* spp. respectively |
| T (generation time) | 1 year | (14) |
| $\gamma$ (strength of selection) | 0.578 (mean across rounds); 0.463, 0.606, 0.590 (rounds 1-3) | Experimental results (*i.e.*, difference in larval survival between treatments) across all rounds, and for each round individually |
| $h^2$ (heritability) | 0.370, 0.228 | Experimental results; GCTA approach. Values are for the LD-pruned, and unpruned SNP data, respectively |
| $\sigma_p^2$ (phenotypic variance) | 0.278, 0.258 | Experimental results; GCTA approach. Values are for the LD-pruned, and unpruned SNP data, respectively |

**Table S9**. Metrics of warming used in comparison with our estimated maximum rate of evolutionary change. Rates of warming are estimated between 2020 and 2050 for metrics 1,4 and 5; between 1976-2005 and 2021-2050 for metrics 2 and 3; and between 1981 and 2022 for metric 6. Results for warming metric 1 are shown in Figure 4 of the main text, while results for 2-6 are shown in Supplemental Figure S7.

| | **Description** | **Rate (°C/year)** | **Data source** |
|---|---|---|---|
| 1 | Rate of change in mean daily temperature in the spring across the southern portion of the *Ae. sierrensis* distribution under a moderate warming scenario (RCP 4.5) | 0.0256 | CHELSA (36) |
| 2 | Rate of change in annual mean temperature across the southeastern U.S. under RCP 4.5 | 0.031 | U.S. Global Change Research Program (37) |
| 3 | Rate of change in annual mean temperature across the southeastern U.S. under an upper warming scenario (RCP 8.5) | 0.035 | U.S. Global Change Research Program |
| 4 | Rate of change in maximum daily temperature in the spring across the southern portion of the *Ae. sierrensis* distribution under RCP 4.5 | 0.0396 | CHELSA |
| 5 | Rate of change in annual mean temperature across the southern portion of the *Ae. sierrensis* distribution under RCP 4.5 | 0.0154 | California Basin Characterization Model (38) |
| 6 | Recently observed rate of change in annual mean temperature across western North America | 0.027 | NOAA |
